# Divergent Proteomic Responses to Lead Exposure in House Sparrows (*Passer domesticus*)

**DOI:** 10.64898/2026.11.10.765757

**Authors:** Zeshan Ali, Max Mclennan-Gillings, Yuheng Sun, Martin Longford, Ondi L. Crino, Mark P. Taylor, Simon C. Griffith, Paul A. Haynes

## Abstract

Chronic exposure to environmental contaminants can drive population decline but also exert selective pressure favouring adaptive molecular responses that mitigate toxicity. Australian lead (Pb) mining and smelting towns provide a natural experiment to study such contemporary adaptation in the house sparrow (*Passer domesticus*), an invasive commensal species that has persisted in heavily contaminated environments for over a century. The Broken Hill population is of particular interest as it has remained geographically isolated with limited gene flow, whereas the Port Pirie population is connected to surrounding regions. Here, we used data-independent acquisition mass spectrometry to compare the blood and liver proteomes of wild-caught sparrows from both towns and identify divergent molecular responses to lead. Lead exposure altered the mitochondrial proteome and disrupted RNA splicing in both populations, but the two towns diverged sharply in haem biosynthesis and protein quality control. We identified a mechanistic cascade in which lead inhibits ALAD, depleting intracellular haem, stabilising the transcriptional repressor BACH1, and suppressing globin expression. Broken Hill birds maintained ALAD abundance, upregulated proteasomal subunits (PSMC) and HSP70 family chaperones, and preserved significantly higher haemoglobin concentrations despite equivalent blood lead levels. Port Pirie birds exhibited ALAD collapse, BACH1 accumulation, and reliance on autophagy rather than active protein quality control. Liver proteomics confirmed tissue-level convergence of the haem pathway disruption. Together, these findings reveal that over a century of isolation under sustained lead exposure has shaped population-specific molecular strategies in Broken Hill sparrows, with enhanced proteostasis and preserved haem biosynthesis representing signatures of local adaptation.

## 1. Introduction

Lead (Pb) is a persistent environmental contaminant with widespread anthropogenic sources and substantial impacts on humans and wildlife [1–8]. Although regulatory interventions have reduced human exposure, lead contamination remains a significant threat to wildlife, particularly in mining and smelting environments [9]. Birds are especially susceptible through ingestion of contaminated soil and food, as well as lead fragments from ammunition [10–12]. Chronic sublethal exposure can impair fitness-related traits, including reproduction and survival, without causing overt mortality, potentially contributing to population-level effects [13, 14].

Persistent environmental contaminants may also impose selection on exposed populations. Where exposure generates consistent and heritable fitness costs, natural selection can favour traits that increase tolerance, potentially leading to local adaptation. However, anthropogenic contaminants are often relatively recent environmental stressors, and the rate of environmental change may exceed the capacity of populations to adapt [15]. Nevertheless, rapid adaptive responses to anthropogenic contaminants have been documented, including responses to polychlorinated biphenyls in Atlantic tomcod, pollution in killifish, and copper in *Drosophila melanogaster* [16, 17]. Understanding the molecular basis of such responses is therefore important for determining how wildlife populations respond to persistent environmental contamination.

The house sparrow (*Passer domesticus*) provides a useful system for investigating adaptation to chronic lead contamination. Introduced into Australia in the 1800s, house sparrows have established populations in the mining and smelting towns of Broken Hill, New South Wales, and Port Pirie, South Australia, where more than a century of mining and smelting has produced strong spatial gradients in environmental lead exposure [11, 18], providing an opportunity to investigate population-specific responses to contrasting contamination histories. House sparrows are relatively sedentary and exhibit moderate population structure and limited gene flow [19, 20]. The Port Pirie population is geographically connected to the broader South Australian population, including the presumed source or parent population around Adelaide, and therefore provides a useful reference for comparison with the more isolated Broken Hill population. In contrast, approximately 100 years of geographical isolation of the Broken Hill population has substantially reduced gene flow with neighbouring populations, creating conditions in which selection associated with chronic lead exposure may be more readily detectable [18–20]. This contrasting demographic history provides a useful context for investigating local adaptation to chronic lead exposure, as greater isolation at Broken Hill may facilitate the accumulation and detection of population-specific adaptive variation, while ongoing connectivity at Port Pirie may constrain such differentiation.

Previous genomic analyses of Australian house sparrows identified signatures of selection in populations from mining and smelting environments, providing evidence for population differentiation associated with environmental exposure [21]. However, genomic variation alone does not reveal how these differences translate into molecular and physiological phenotypes. Changes in gene expression and protein abundance provide complementary information on the functional consequences of environmental exposure and may contribute directly to phenotypic divergence between populations [16, 22].

Here, we used quantitative mass spectrometry to characterise proteomic responses to chronic lead exposure in blood and liver from house sparrows in Broken Hill and Port Pirie. Proteomics provides a functionally proximal measure of cellular state because proteins directly mediate metabolic, regulatory, and stress-response processes [23]. Analysis of both tissues further distinguishes systemic responses captured in blood from hepatic responses associated with detoxification and xenobiotic metabolism [24, 25]. Because protein abundance can be regulated independently of transcript abundance through post-transcriptional processes and protein turnover, proteomics also provides information that cannot be inferred directly from transcriptomic measurements [26]

We therefore aimed to determine whether house sparrows from two geographically distinct, chronically contaminated populations exhibit divergent proteomic responses to lead exposure. Untargeted quantitative proteomics was performed on blood and liver from high- and low-lead birds at each site, with exposure groups defined by blood lead concentration and sampling distance from the primary contamination source.

## 2. Materials and Methods

### 2.1. Sampling locations

Samples were collected from two towns in Australia: Broken Hill, New South Wales, and Port Pirie, South Australia (Figure 1). Broken Hill has one of the largest silver-lead-zinc orebodies in the world, and the town is built around the active mine into two residential districts. Mining activities have been ongoing since 1885, creating a soil lead gradient in town, with higher lead levels near the mine. The ore from the mine is transported to Port Pirie in South Australia for smelting, a distance of ∼400 km. The smelter was built in 1889, and, like the mine in Broken Hill, has left a gradient of lead in the soil. The sampling sites in this study were selected based on soil lead contamination and blood lead levels (BLL) from previous studies [11, 27–29] and were divided into four groups: Broken Hill high lead (BH High), Broken Hill low lead (BH Low), Port Pirie high lead (PP High), and Port Pirie low lead (PP Low). Blood samples were collected from sparrows captured in 3 locations in Broken Hill (2 with high BLLs and 1 with a low BLL) and 5 locations in Port Pirie (4 with high BLLs and 1 with a low BLL).

**Figure 1:**
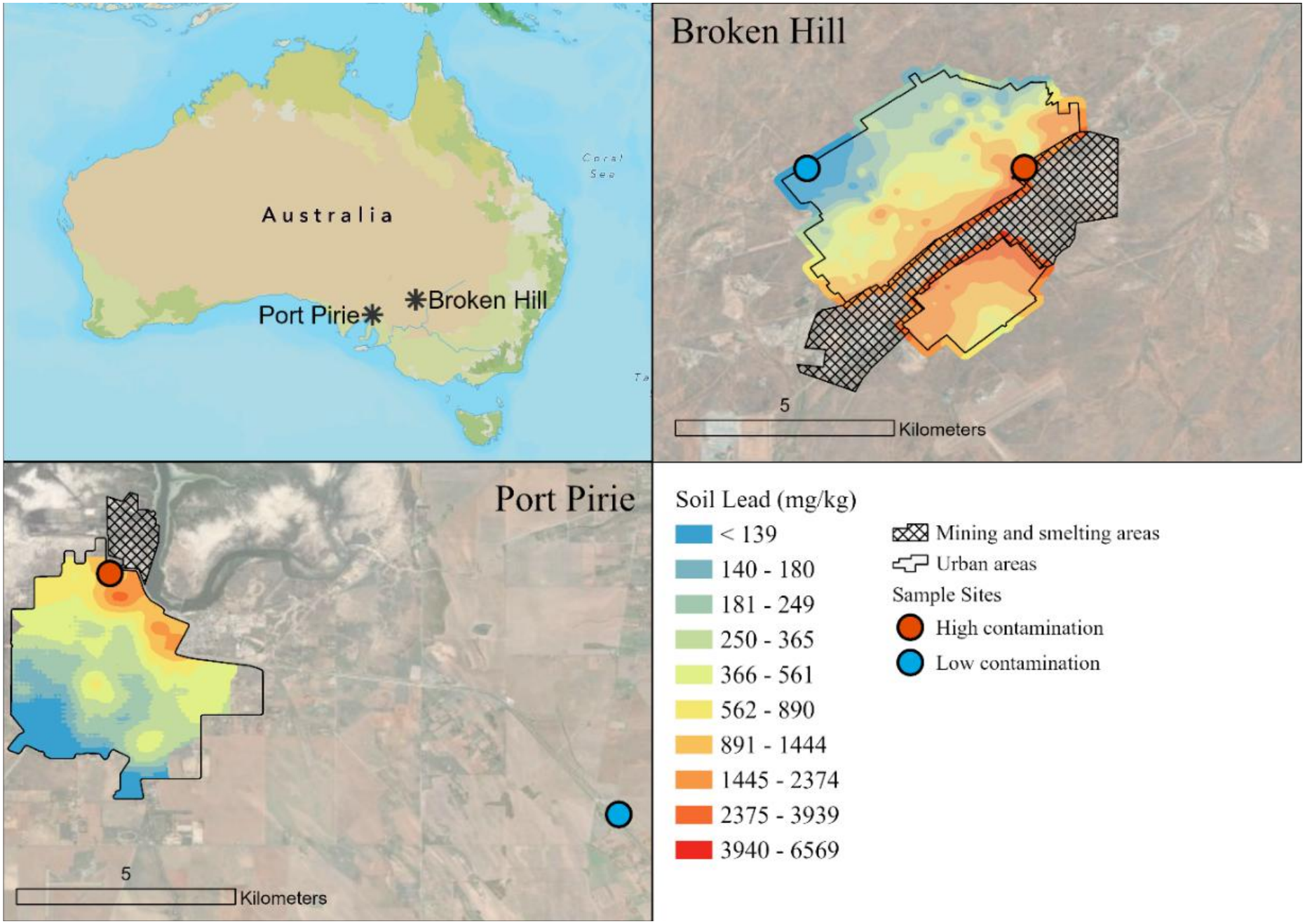
Map of Broken Hill (b) and Port Pirie (c) sampling sites.

### 2.2. Soil trace metal analysis

At each site, surface soils (0–2 cm depth) were collected in triplicate and analysed for metal concentrations following established protocols [11, 29]. Samples were sieved to <250 µm, transferred to Mylar film analysis cups, and analysed for metal concentrations (mg/kg) using X-ray fluorescence (XRF) spectrometry with a Vanta portable XRF spectrometer (Olympus-IMS). The spatial maps in Figure 1b were generated using soil lead data collected during the 2023 field season as part of a previously published study [15], supplemented with additional measurements from the current study. For Port Pirie, site-specific soil lead measurements were complemented with surface-soil data from a 2013 roadside survey conducted on a regular grid across the town. We report data on As, Cr, Cu, Fe, Mn, Ni, Pb, Ti, V, and Zn, although we focus on Pb as the primary contaminant of concern associated with mining and smelting activities in this region.

### 2.3. Biological sampling

Approval for bird capture and sampling was granted by the Macquarie University Animal Ethics Committee (AEC#2020/011-14). Sparrows were captured using mist nets, and captured birds were banded and identified for age (juvenile/adult) and sex (female/male). Body mass, wing chord, and tarsus length were measured for each individual. Blood samples were obtained by puncturing the brachial vein with a 26-gauge hypodermic needle. BLLs were measured using a Meridian Bioscience LeadCare Plus blood Pb analyser, which employs anodic stripping voltammetry (ASV) to quantify lead in whole-blood samples mixed with a dilute HCl solution. The instrument has a manufacturer-specified detection range of 1.9–65.0 µg/dL for human blood. For avian blood, ASV underestimates lead levels and was corrected as described by Gillings *et al.* [15], shifting the ASV-detectable blood lead range from the proprietary 1.9–65 μg/dL to 5.97– 111.0 μg/dL. Blood haemoglobin (Hb) levels (g/L) were measured using a HemoCue Hb 201+ instrument, using methods described previously [11].

For proteomic analysis, a whole blood sample of ∼20-30 μL was collected into an EDTA tube, and 15 μL of blood was transferred to a microtube. A subset of captured adult male birds was euthanised, and liver and brain samples were collected. All samples were frozen in a dry shipper, transported in a freezer at -20 °C, and then transferred to a freezer at -80 °C for long-term storage. Because of the balanced design for males and females, sex was not considered a covariate and was not controlled for.

### 2.4. Whole blood proteome sample preparation

Whole blood samples (n = 60, 15 from each group) were prepared using the Mitra VAMS (Volumetric Absorptive Microsampling) dried blood method described by Molloy *et al.* [30], with certain modifications specific to avian blood. Samples were evenly split between the two localities, with 30 birds from Broken Hill and 30 from Port Pirie. The four groups comprised two Broken Hill groups, each drawn from a single site (high-lead, Argent Street, n = 15; low-lead, Brookfield Avenue, n = 15), and two Port Pirie groups (high-lead, spanning George Street [n = 10], Hallam Street [n = 2], Warnertown Road [n = 2], and Senate Road [n = 1]; low-lead, Augusta Highway, n

= 15). Blood samples (15 μL) were thawed on ice and 15 μL of the whole blood was diluted with 45 μL of phosphate-buffered saline (PBS). Diluted blood samples were transferred to a 96-well U-bottom plate. VAMS devices were placed on the samples until fully saturated, and the samples were then dried at room temperature for 20 min. The devices were transferred to specimen foil ziplock bags containing desiccant and incubated at 4 °C for 3 h. Following incubation, devices were removed from the specimen bags, their tips were detached from the spindles, and the tips were placed into a 96-well deep-well filter plate. Tips were washed with washing buffer (800 μL; 500 mM LiCl, 100 mM Tris) at 300 rpm for 24 h. The washing solution was removed by centrifuging for 4 min at 2,000 g, and the tips were further washed three times with 800 μL of washing buffer for 4 min at 700 rpm each, followed by centrifugation at 2,000 g for 4 min after each wash. Following the final wash, dried tips were transferred to a 96-well plate, and 130 μL of lysis buffer was added (1% sodium deoxycholate, 100 mM triethylammonium bicarbonate, 10 mM tris(2-carboxyethyl) phosphine, 40 mM chloroacetamide). The plate was heated to 95 °C for 10 min. Upon cooling, 1 μg of trypsin (Trypsin Gold, mass spectrometry grade; Promega, Madison, Wisconsin, USA) was added to each well, and samples were digested at 37 °C for 18 h. The trypsin reaction was quenched by adding 200 μL of 0.5% formic acid. Tips were removed from the digest using forceps, and the plate was centrifuged at 2,000 g to pellet sodium deoxycholate. The supernatant was desalted using an Oasis HLB 96-well plate (10 mg sorbent; Waters, Milford, MA, USA) as described previously [31]. Quality control (QC) samples were prepared by pooling equal volumes of diluted whole blood from randomly selected study samples. The pooled material was divided into six aliquots, prepared alongside the biological samples, and distributed randomly across the plate to monitor technical variability throughout the acquisition.

A second cohort (n = 23), hereafter referred to as the secondary cohort, was prepared independently using the same workflow, with whole blood diluted 10-fold rather than 3-fold to accommodate smaller sample volumes. This cohort comprised 8 birds from Broken Hill (high-lead, Argent Street, n = 4; low-lead, Brookfield Avenue, n = 4) and 15 from Port Pirie (high-lead, spanning George Street [n = 1], Warnertown Road [n = 6], and Senate Road [n = 1]; low-lead, Augusta Highway, n = 7). All steps were identical to those in the main cohort except for desalting, which was performed using an ethyl acetate phase-transfer stage-tip method [32]. Briefly, digested peptide solutions were mixed with an equal volume of 99% ethyl acetate/ 1 % TFA, and the aqueous phase was loaded onto equilibrated stage-tips. Tips were washed twice with 99% ethyl acetate / 1% TFA and once with 0.1% TFA, then eluted with 80% acetonitrile / 5% ammonium hydroxide. Eluates were vacuum-dried and reconstituted in 0.5% formic acid to a final concentration of 0.1 µg/µL.

### 2.5. Liver proteome sample preparation

Liver samples (n = 24) were collected from a subset of birds across both localities, comprising 11 from Broken Hill (high-lead, Argent Street, n = 6; low-lead, Brookfield Avenue, n = 5) and 14 from Port Pirie (high-lead, collected from George Street [n = 5], and Warnertown Road [n = 2], low-lead, Augusta Highway, n = 6). Liver tissue was homogenised by bead beating (glass beads, 710–1,180 µm) in 1 mL ice-cold lysis buffer (50 mM Tris-HCl, pH 7.5, 150 mM NaCl, 1% NP-40, 1 mM EDTA, 0.1% SDS, 1% v/v protease inhibitor cocktail) followed by probe sonication for 15 seconds. Insoluble debris was removed by centrifugation at 13,000 × g for 15 minutes at 4°C, and 50 µL aliquots of clarified supernatant were retained for processing. Proteins were reduced with 20 mM DTT (30 minutes, room temperature), alkylated with 50 mM IAA (30 minutes, dark), and quenched with 40 mM DTT. Proteins were precipitated by methanol– chloroform extraction, air-dried, and reconstituted in 8 M urea / 100 mM Tris-HCl (pH 8.5) overnight at 4°C. Protein concentration was determined by BCA assay, and 50 µg aliquots were diluted tenfold into 100 mM Tris-HCl (pH 8.5) and digested with trypsin (50:1, w/w) for 16 hours at 37°C. Digestion was quenched with 99% ethyl acetate / 1% TFA, and peptides were desalted using SDB-RPS stage-tips, eluted with 80% acetonitrile / 5% ammonium hydroxide, vacuum-dried, and reconstituted in 0.5% formic acid to a final concentration of 0.2 µg/µL.

### 2.6. HL-60 cell line lead treatment

HL-60 cells were purchased from ATCC (HL-60, CCL-240 ™) and cultured in RPMI 1640 medium supplemented with 10% fetal bovine serum and 1% penicillin-streptomycin. Cells were passaged when they reached approximately 85% confluency, and cell viability was assessed using Trypan Blue. Approximately 10^6^ cells were seeded into 6-well plates (5 control, 5 treated). Treated wells received 100 µg/dL lead acetate for 24 hours; control wells received vehicle only. Following treatment, cells were harvested, washed with DPBS and lysed in 4% sodium deoxycholate (SDC) in 100 mM Tris-HCl (pH 8.5) by probe sonication. Protein concentration was determined by BCA assay. Proteins were reduced with TCEP (10 mM) and alkylated with chloroacetamide (40 mM) in 4% SDC buffer at 95 °C for 10 minutes. After cooling, samples were diluted to a final concentration of 1% SDC and digested with sequencing-grade trypsin (Promega, Madison, Wisconsin, USA) at a 1:50 enzyme-to-substrate ratio. Peptides were desalted using the ethyl acetate phase-transfer-based stage tip method as described above for liver tissue. Peptide concentrations were determined using a Nanodrop (Thermo Fisher Scientific, CA, USA) and standardized to 0.1 μg/uL in 0.5% formic acid.

### 2.7. Blood and liver LC-MS/MS single injection

Tryptic peptides (450 ng) from whole blood and liver were analyzed on a Vanquish Neo UHPLC system (Thermo Scientific, CA, USA) coupled to an Orbitrap Exploris-480 mass spectrometer (Thermo Scientific, CA, USA). Peptides were loaded onto an Acclaim PepMap 100 C18 trap column (5 μm particle size, 100 Å pore size, 300 μm × 5 mm; Thermo Fisher Scientific) and separated on a 300 mm C18 analytical column (75 μm internal diameter) packed with Dr. Maisch Reprosil-Pur resin (3 μm particle size, 120 Å pore size). Both trap and analytical columns were maintained at 45 °C. Peptides were loaded in buffer A (0.1% formic acid) and eluted using buffer B (80% acetonitrile, 0.1% formic acid) with the following gradient profile: 2% to 37.5% buffer B over 60 min (linear gradient), 37.5% to 95% buffer B over 3 min, and held at 95% buffer B for 10 min. Peptides were electro-sprayed using a stainless-steel emitter (Nano Bore O.D. 150 um, I.D. 30 um, Thermo Scientific, CA, USA). Data were acquired in data-independent acquisition (DIA) mode. MS1 scans were acquired at a resolution of 120,000 for blood samples and 60,000 for liver samples, with a scan range of 350–1,450 m/z. Maximum injection time was set to automatic, with automatic gain control set to standard. Each MS1 scan was followed by 20 MS2 scans using variable overlapping wide windows spanning 350–1,450 m/z. Precursor ions were fragmented using higher-energy collisional dissociation (HCD) at 27% normalized collision energy, and fragment ions were analysed in the Orbitrap at 30,000 resolution.

### 2.8. LC-MS for multi-injection gas-phase fractionation (GPF)

A separate empirical spectral library was generated for the blood proteome using the same LC-MS system described above. Desalted peptide samples from whole blood were pooled and analyzed using a multi-injection gas-phase fractionation (GPF) strategy. The pooled sample was injected seven times to cover a precursor m/z range of 395–1,450. The m/z ranges covered by each sequential injection were: 395–505, 495–605, 595–705, 695–805, 795–905, 895–1,005, and 1,005–1,450. Chromatographic separation conditions were identical to those used for single-injection analysis. MS1 spectra were acquired at 60,000 resolution, and precursor ions were fragmented using higher-energy collisional dissociation (HCD) at 27% normalized collision energy. Automatic gain control was enabled with a normalized AGC target of 1,000 and automatic maximum injection time. Each MS1 scan was followed by 51 staggered MS2 scans. For the first six injections, 4 m/z isolation windows were used, while the final injection (1,005–1,450 m/z) employed 51 isolation windows of 28 m/z to cover the high-mass precursor range. MS2 spectra were acquired at a resolution of 30,000.

### 2.9. HL-60 and whole blood secondary cohort LC-MS/MS single injection

Peptides derived from the HL-60 cell line experiment and whole blood secondary cohort were analysed using a Vanquish Neo UHPLC system (Thermo Fisher Scientific, CA, USA) coupled to an Orbitrap Astral mass spectrometer (Thermo Fisher Scientific, CA, USA). Peptide separation was performed on a nanoEase M/Z HSS T3 C18 column (1.8 μm particle size, 100 Å pore size, 300 μm × 150 mm; Waters, Milford, MA, USA) maintained at 45 °C. Peptides were loaded in mobile phase A (0.1% formic acid in water) and separated using mobile phase B (80% acetonitrile, 0.1% formic acid) at a flow rate of 5 μL/min. The chromatographic gradient consisted of a linear increase from 2% to 35% mobile phase B over 17 min, followed by an increase to 95% mobile phase B over 1 min, and a 6 min column wash at 95% mobile phase B. Data were acquired in data-independent acquisition (DIA) mode. Full MS (MS1) scans were acquired in the Orbitrap mass analyser at a resolution of 240,000 (at m/z 200) across an m/z range of 380–980, with the absolute automatic gain control (AGC) target set to 5 × 10⁶ ions. Precursor ions were fragmented using higher-energy collisional dissociation (HCD) with a normalized collision energy (NCE) of 25. MS/MS spectra were acquired in the Astral mass analyser using 2 Da isolation windows across the precursor m/z range of 380–980. A total of 300 DIA scan events were collected per acquisition cycle. Fragment ions were detected over an m/z range of 150–2000, with a maximum injection time of 5 ms and an absolute AGC target of 2 × 10⁴ ions.

### 2.10. Spectral Processing

Thermo RAW files from blood analysis were processed using the DIA-NN computational platform (v2.2.0) [33] through a three-step DIA spectral analysis workflow. A previously published high-quality *Passer domesticus* protein database (14,260 sequences) was used for spectral matching [34, 35]. The database contained custom headers and lacked protein and gene annotations. To address this, the database was mapped and annotated using eggNOG-mapper [36], and the headers were subsequently formatted to UniProt-style entries using custom R scripts.

When a gene or protein name mapped to multiple sequences, the corresponding annotation was excluded to avoid ambiguity, and the original custom identifier was retained. Consequently, each protein sequence retained a unique identifier, and no two sequences shared the same protein or gene name. Sequences without annotation matches retained their original custom identifiers.

In the first step, this curated FASTA file was used to generate a predicted *in silico* library using DIA-NN neural networks. MS2 mass accuracies were set to 10 ppm, match-between-runs (MBR) was enabled, and the following parameters were applied for *in silico* library generation: trypsin as the protease with one missed cleavage allowed; fixed modifications included carbamidomethylation of cysteine and N-terminal methionine excision; no variable modifications were selected (to limit the search space); peptide length of 7–30 residues; precursor charge range of 2–4; precursor m/z range of 350–1,450; and fragment m/z range of 200–1,800. In the second step, the *in-silico* library was used to search the multi-injection gas-phase fractionation (GPF) files for each sample type. MS1 scan windows were optimized automatically by DIA-NN. Precursors were filtered at 1% false discovery rate (FDR), and proteotypicity was specified using gene names from the database. This process generated sample-specific empirical spectral libraries. In the third step, these empirical libraries were used to search single-injection DIA files with identical parameters, except that MBR was disabled. Quantification was performed using the QuantUMS (high-precision) strategy [37] with cross-run normalization.

For the liver whole-blood secondary cohort and the HL-60 cell line, no GPF library was prepared, and the data were processed using the in-silico library. All settings were kept the same, except for two variations. The datasets were searched with MBR enabled, and the precursor range for the Astral data was 380-980 m/z, with 5 ppm mass accuracy for MS1. The HL-60 cell line data were searched against the human protein database, downloaded from Uniprot and containing 20,436 reviewed sequences. All datasets were searched separately.

### 2.11. Differential abundance analysis

All proteomics data analyses were performed in R (version 4.5.2) using RStudio, unless otherwise specified. Protein abundance analysis was performed using the peptide-level quantification model MS-EmpiRe [38] implemented in the MS-DAP R package (version 1.2.2) [39] with the following settings: minimum detections = 2, minimum quantifications = 2, minimum peptides per protein = 2, filter by contrast = TRUE, normalization algorithm = variance stabilizing normalization (VSN) with between-protein mode. MS-EmpiRe was used as the differential expression algorithm with a q-value threshold of 0.05 and a log₂ fold-change cut-off of 0.5. Proteins initially identified as significant by the MS-EmpiRe model were further filtered using the DEqMS and MSqRob models and were considered significant if the p-value in both models was < 0.05. Because the sexes were approximately balanced across the four groups (31 males, 29 females), sex was not treated as a covariate and was not controlled for in downstream analyses.

For HL-60 cell line data, MSqRob in MSDAP was used for the differential analysis with a q-value threshold of 0.05 and a log₂ fold-change cut-off of 0.5.

### 2.12. Proteome Divergence Response Analysis

The protein MaxLFQ intensity matrix was generated from the primary ‘report.parquet’ file using the DIA-NN R package with the following quality thresholds: Q.Value ≤ 0.01, PG.Q.Value ≤ 0.05, Global.Q.Value ≤ 0.01, and Global.PG.Q.Value ≤ 0.01. The resulting protein group dataset was filtered in two steps. First, protein groups identified by fewer than two peptides were removed. Second, protein groups with more than 40% missing values across all samples were excluded, retaining only proteins with at least 60% valid quantification values for downstream analysis. Protein intensities were log₂-transformed, and missing values were imputed by random sampling from a normal distribution with a mean downshifted by 1.8 standard deviations (SD) and a width of 0.3 SD relative to the abundance distribution of all proteins within each sample [40]. The final matrix contained 1,292 proteins and 57 samples. Each group contained 15 samples except PP LL, which contained 12 samples. Linear models were fitted to each protein using the limma package [41]. The design matrix encoded a full factorial model:

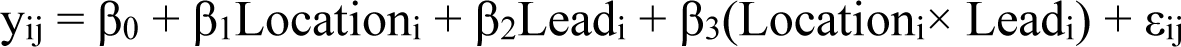

where y_ij_ represents the outcome for individual j in group i. Location was coded with BH as the reference and Lead with Low (low lead) as the reference. The interaction coefficient β_3_ quantifies the difference in lead response between PP and BH birds: a positive value indicates a stronger, more positive response in PP than in BH. Empirical Bayes moderation was applied via the eBayes function, which estimates a prior variance (S₀^2^) and prior degrees of freedom (d0) from the distribution of residual variances across all proteins. Each protein’s residual variance is then shrunk toward this prior, producing moderated t-statistics with increased posterior degrees of freedom. Four contrasts were tested per protein: (i) the interaction term (β3), testing whether the lead response differs between locations; (ii) the BH-specific lead effect (β2), since BH is the reference level; (iii) the PP-specific lead effect (β2 + β3), extracted using contrasts.fit; and (iv) the population baseline difference (β1), comparing BH Low versus PP Low. P-values were corrected for multiple testing using the Benjamini-Hochberg procedure to control the false discovery rate (FDR). For each protein, the per-population lead response was also computed as the difference in group means: BH response = mean(BH High) minus mean(BH Low), and PP response = mean(PP High) minus mean(PP Low). These values were used for visualisation and for classifying the response pattern of each protein. Patterns were assigned as follows: Opposite (both populations respond with absolute log2FC above 0.15 but in different directions), BH stronger or PP stronger (same direction but one population’s response exceeds 1.5 times the other), BH only or PP only (one population responds above 0.15 while the other remains below 0.1), or Similar (responses do not meet any divergence criterion).

Protein quality control (PQC) pathway analysis was performed by assigning proteins to functional categories (i.e. Proteasome, Ubiquitin, Chaperone, Autophagy, ER/UPR etc.) based on keyword matching against gene symbols and functional descriptions from eggNOG-mapper annotations. For each category, the mean BH and PP lead responses were computed across all member proteins to assess population-level deployment of each proteostasis arm.

### 2.13. Gene Ontology and Pathway Enrichment analysis

Gene ontology (GO) enrichment analysis was performed using the clusterProfiler package in R [42]. Sparrow protein group identifiers were first mapped to gene symbols using eggNOG-mapper annotations. Because GO and KEGG pathway databases are not available for *Passer domesticus*, gene symbols were mapped to human orthologs using the org.Hs.eg.db annotation database. Enrichment was tested separately for three GO domains (Biological Process, Molecular Function, Cellular Component) and for KEGG pathways. For KEGG analysis, gene symbols were converted to Entrez IDs using the bitr function. P-values were adjusted for multiple testing using the Benjamini-Hochberg method and enrichment was considered significant at an adjusted p-value below 0.05.

### 2.14. Lead-Heme statistical modeling

The relationship between blood lead and haemoglobin was modelled using quantile regression at the conditional median (τ = 0.5), implemented in R using the quantreg package, as residuals from the Broken Hill ordinary least squares model violated the normality assumption (Shapiro-Wilk test). Blood lead was natural log-transformed before modelling, to linearise the dose-response relationship. Sex was not included as a covariate because haemoglobin did not differ significantly between males and females (Δ = 2.1 g/L; Welch t-test: p = 0.562), and a preliminary model including sex produced a non-significant sex coefficient (β = −0.68 g/L, p = 0.841) with negligible change to the blood lead slope (Δβ = 0.09 g/L).

Six models were fitted: a combined model testing a location × log (blood lead) interaction (A1); town-stratified continuous models for Broken Hill (A2) and Port Pirie (A3); town-stratified categorical models comparing Low versus High exposure groups within each town (A4, A5); and a pooled model across both towns without a location term (A0). Standard errors and p-values were estimated via bootstrap resampling of (x, y) pairs (R = 2,000), as the default sparsity-based estimator performs poorly at small sample sizes when predictor values are tied due to detection-limit substitution in the low-exposure groups. Model fit was summarised using the Koenker-Machado pseudo-R². Spearman’s rank correlations were computed as a non-parametric complement to each continuous model.

### 2.15. Citrate Synthase Activity Assay

Citrate synthase (CS) activity was measured using the protocol defined by Nord *et al.* [43] with minor modifications. Whole blood samples (n = 60) were assigned to randomised blocks of four samples, with each block containing one representative from each group (BH HL, BH LL, PP HL, PP LL), distributed across three 96-well plates. Blood was homogenised in homogenisation buffer consisting of 100 mmol L⁻¹ KH₂PO₄ (pH 7.2), 1 mmol L⁻¹ EGTA, 1 mmol L⁻¹ EDTA, and 1 mmol L⁻¹ phenylmethylsulfonyl fluoride. Homogenates were centrifuged at 1000 × g for 10 min at 4°C, and supernatants were collected. Protein concentration in each supernatant was quantified in duplicate using the Thermo Scientific Pierce BCA Protein Assay Kit.

CS activity was measured in triplicate at 412 nm using a BMG FLUOstar Omega microplate reader maintained at 41°C. Each 200 µL reaction well contained 100 mmol L⁻¹ KH₂PO₄ (pH 7.2), 0.15 mmol L⁻¹ acetyl-CoA, 0.15 mmol L⁻¹ 5,5′-dithiobis-2-nitrobenzoic acid (DTNB), and 0.5 mmol L⁻¹ oxaloacetate (omitted in blanks). The mean slope of the linear phase (ΔA₄₁₂ min⁻¹) was calculated after subtracting the corresponding blank well, and triplicates with a coefficient of variation exceeding 20% or a negative slope were excluded from further analysis. CS activity was calculated as:

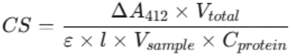

where Δ*A*₄₁₂ is the blank-corrected mean slope (AU min⁻¹), *V*_total_ = 0.200 mL is the total well volume, ε = 14.15 mM⁻¹ cm⁻¹ is the molar extinction coefficient of TNB at 412 nm, *l* = 0.60 cm is the theoretical path length for 200 µL in a 96-well plate, *V*_sample_ is the volume of homogenate loaded (mL), and *C*_protein_ is the protein concentration of the homogenate (µg µL⁻¹, equivalent to mg mL⁻¹). CS activity is expressed as µmol min⁻¹ mg protein⁻¹.

Group differences in CS activity were assessed using Welch’s t-test (assuming unequal variances). Five pairwise comparisons were performed: (i) overall High versus Low lead (pooled across locations), (ii) BH Low versus BH High, (iii) PP Low versus PP High, (iv) BH Low versus PP Low (population baseline), and (v) BH High versus PP High.

### 2.16. Blood metabolite derivatisation

Whole blood samples (2 µL) were randomised within a blocked experimental design and divided into three preparation batches of 24 samples each. Frozen blood aliquots were thawed on wet ice and diluted in 200 µL of 50% methanol in water (v/v). The diluted samples were filtered through a 10 kDa molecular weight cutoff membrane (Millipore, 2 mL or 15 mL devices, regenerated cellulose) by centrifugation (batch 1: 8000 × g for 1 hour; batches 2 and 3: 4000 × g for 2 hours, 4°C) to remove haemoglobin and other high molecular weight proteins. The filtrates were dried in a vacuum centrifuge and reconstituted in 10 µL of 50% methanol. Reconstituted extracts were derivatised with 3-nitrophenylhydrazine (3-NPH), 1-ethyl-3-(3-dimethylaminopropyl) carbodiimide (EDC) and pyridine prepared in 50:50 acetonitrile/water (v/v). 3-NPH was prepared at 200 mM and the EDC/pyridine solution at 120 mM EDC containing 6% pyridine (v/v). Derivatisation was carried out at 40°C for 30 minutes. Under these conditions, 3-NPH reacts with the C4 ketone of 5-aminolevulinic acid via spontaneous hydrazone formation and with carboxylic acid groups via EDC-mediated acylhydrazide coupling, while pyridine serves as a base catalyst. A pooled quality-control sample prepared from a representative subset of biological samples was processed identically with each batch.

### 2.17. Targeted quantification of 5-aminolevulinic acid and porphobilinogen in whole blood

Derivatised samples were analysed on a Shimadzu LCMS-8060NX triple quadrupole mass spectrometer equipped with a Nexera XR SIL-40C X3 autosampler and SCL-40 system controller at the Australian Genome Foundry, Macquarie University. Chromatographic separation was performed on a Shimadzu Shim-pack Scepter C18-120 column (3 µm, 2.1 × 100 mm) maintained at 50°C with an injection volume of 2 µL. Mobile phase A consisted of water with 2 mM ammonium formate and 0.1% formic acid, and mobile phase B consisted of methanol with 2 mM ammonium formate and 0.1% formic acid, delivered at a flow rate of 0.2 mL/min. The gradient program was as follows: 5% B held from 0 to 1.0 min, ramped linearly to 30% B at 3.0 min, held at 30% B until 4.0 min, ramped to 95% B at 7.0 min, held at 95% B until 11.0 min, returned to 5% B at 11.5 min and re-equilibrated until 14.5 min.

The mass spectrometer was operated in negative electrospray ionisation mode with the following source parameters: interface voltage 4.5 kV, interface temperature 400°C, desolvation temperature 650°C, desolvation line temperature 250°C, heat block temperature 400°C, nebulising gas flow 3 L/min, heating gas flow 15 L/min and drying gas flow 3 L/min. Data were acquired in multiple reaction monitoring (MRM) mode over the full 14.5 min run. Two MRM events were configured.

For 5-aminolevulinic acid mono-NPH derivative (precursor m/z 265.10), three transitions were monitored: 265.10 to 152.15 at a collision energy (CE) of 15 V, 265.10 to 137.10 at CE 20 V and 265.10 to 107.10 at CE 25 V. The m/z 152.15 product ion corresponds to the deprotonated 3-NPH diagnostic fragment ([3-NPH−H]⁻, C₆H₆N₃O₂⁻) and was used as the quantifier transition. The m/z 137.10 product ion arises from direct cleavage of the hydrazone N-N bond and served as the primary qualifier. For porphobilinogen, transitions were monitored for both the mono-NPH derivative (precursor m/z 359.20) and the bis-NPH derivative (precursor m/z 495.20). Five transitions were monitored for the mono-NPH form: 359.20 to 273.10 at CE 25 V, 359.20 to 178.15 at CE 20 V, 359.20 to 150.10 at CE 25 V, 359.20 to 122.05 at CE 25 V and 359.20 to 92.00 at CE 25 V. Five transitions were monitored for the bis-NPH form: 495.20 to 478.10 at CE 20 V, 495.20 to 340.10 at CE 20 V, 495.20 to 299.10 at CE 25 V, 495.20 to 178.15 at CE 25 V and 495.20 to 150.05 at CE 30 V. The mono-NPH form yielded substantially higher signal than the bis-NPH form and was used for quantification, with the m/z 122.05 product ion (nitrophenyl carbanion, C₆H₄NO₂⁻) selected as the quantifier transition based on its consistently high intensity across all samples. The dominance of this low-mass fragment for porphobilinogen is attributed to efficient charge-directed fragmentation facilitated by the electron-rich pyrrole ring of the PBG backbone.

### 2.18. Data processing and batch correction for targeted metabolite data

Peak integration was performed in Shimadzu LabSolutions. Quantifier transition areas were extracted for 5-aminolevulinic acid (265.10 to 152.15) and porphobilinogen (359.20 to 122.05). Product ion agreement was assessed before statistical analysis by computing pairwise Spearman rank correlations and ion-ratio coefficients of variation across all biological samples for each combination of product ions within each analyte. All qualifier transitions showed high agreement with the corresponding quantifier (Spearman’s rho greater than 0.98, ion ratio CV below 7%), confirming that quantification was not confounded by co-eluting interferences.

Batch effects across the three batches were corrected using pool-based normalisation. For each analyte, a correction factor was calculated as the mean pool area across all three batches divided by the pool area of the corresponding batch, and all sample areas within that batch were multiplied by this factor. As a sensitivity analysis, median-centering normalisation was also applied, in which each sample area was divided by the within-batch median and multiplied by the global median. Both approaches yielded concordant statistical results for all tests. The ALA/PBG ratio was calculated from pool-normalised areas for each sample as a functional readout of ALAD enzyme inhibition, where an elevated ratio indicates accumulation of the ALAD substrate (ALA) relative to its product (PBG). For individual analytes and for the ALA/PBG ratio, log₂-transformed pool-normalised values were analysed using a two-way analysis of variance with Location (Broken Hill, Port Pirie), Lead level (Low, High) and their interaction as fixed effects, using Type II sums of squares. Pairwise group comparisons were performed using Tukey’s Honest Significant Difference test with adjustment for multiple comparisons via the emmeans package.

## 3. Results and Discussions

Broken Hill is located in the far western region of New South Wales, approximately 1,100 km from Sydney, at an elevation of 315 m above sea level. The town is characterised by a hot semi-arid climate with a mean annual rainfall of 265 mm. Port Pirie is a coastal town in South Australia situated 223 km north of Adelaide at an elevation of 4 m above sea level, with a mean annual rainfall of 345 mm. The two towns are approximately 400 km apart and share broadly similar semi-arid climatic conditions. The soil metal concentrations are provided in Supplementary Table S1. A summary of biometric data is provided in Table 1 (for all samples from adult individuals, for all individuals including juveniles the data is provided in Supplementary Table S2) and Table 2 (for proteomics samples only).

**Table 1:**
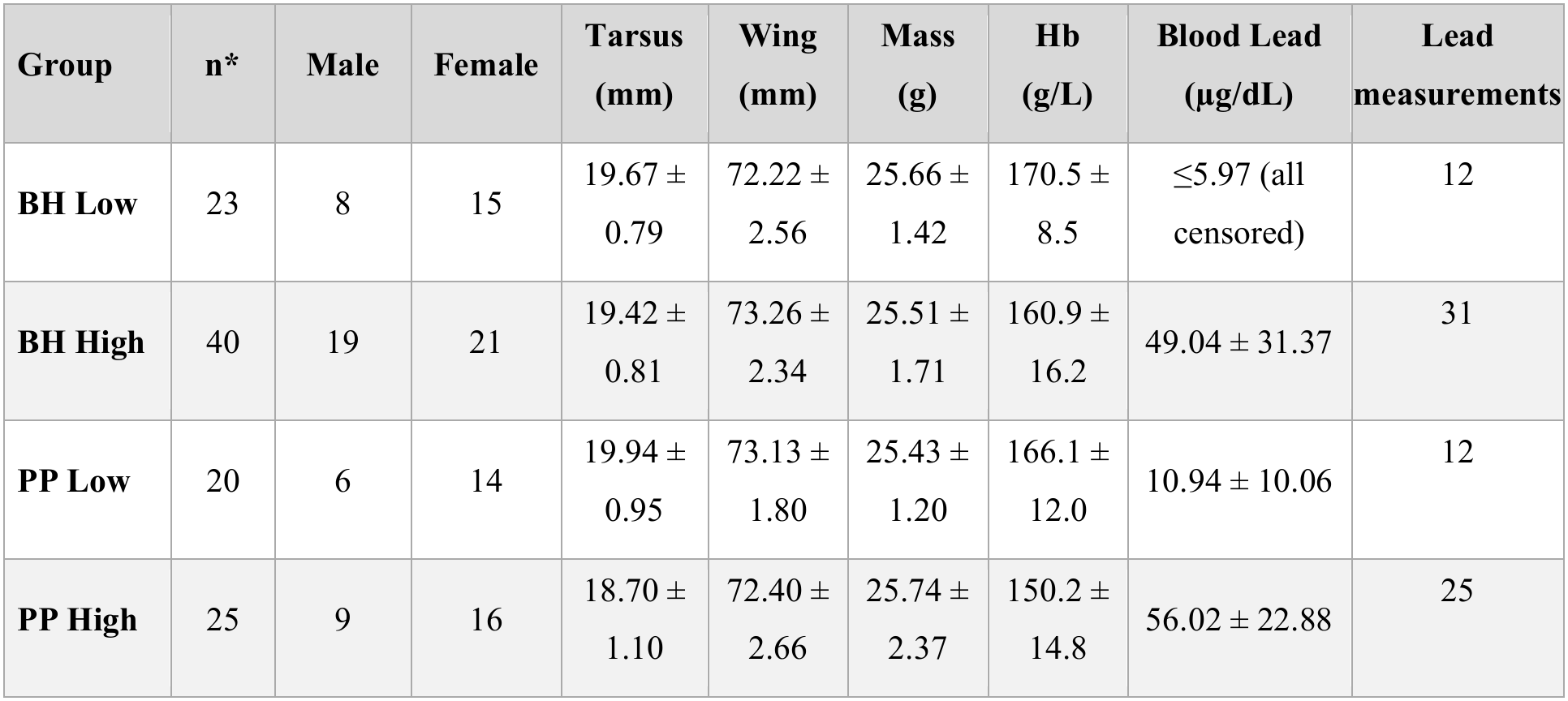
Subject characteristics of house sparrows sampled across Broken Hill and Port Pirie. Values are mean ± SD. *Only adult individuals are included in biometric, blood lead levels (BLL) and haemoglobin summary statistics.

| **Group** | **n\*** | **Male** | **Female** | **Tarsus (mm)** | **Wing (mm)** | **Mass (g)** | **Hb (g/L)** | **Blood Lead (µg/dL)** | **Lead measurements** |
|---|---|---|---|---|---|---|---|---|---|
| **BH Low** | 23 | 8 | 15 | 19.67 ± 0.79 | 72.22 ± 2.56 | 25.66 ± 1.42 | 170.5 ± 8.5 | ≤5.97 (all censored) | 12 |
| **BH High** | 40 | 19 | 21 | 19.42 ± 0.81 | 73.26 ± 2.34 | 25.51 ± 1.71 | 160.9 ± 16.2 | 49.04 ± 31.37 | 31 |
| **PP Low** | 20 | 6 | 14 | 19.94 ± 0.95 | 73.13 ± 1.80 | 25.43 ± 1.20 | 166.1 ± 12.0 | 10.94 ± 10.06 | 12 |
| **PP High** | 25 | 9 | 16 | 18.70 ± 1.10 | 72.40 ± 2.66 | 25.74 ± 2.37 | 150.2 ± 14.8 | 56.02 ± 22.88 | 25 |

**Table 2:** Subject characteristics of the subset used for proteomic analysis. Values are mean ± SD.

| **Group** | **n** | **Male** | **Female** | **Tarsus (mm)** | **Wing (mm)** | **Mass (g)** | **Hb (g/L)** | **Blood Lead (µg/dL)** | **Lead measurements** |
|---|---|---|---|---|---|---|---|---|---|
| **BH Low** | 15 | 8 | 7 | 19.57 ± 0.86 | 72.33 ± 3.02 | 25.72 ± 1.20 | 167.5 ± 6.9 | ≤5.97 (all censored) | 11 |
| **BH High** | 15 | 7 | 8 | 19.51 ± 0.88 | 73.27 ± 2.82 | 24.69 ± 0.69 | 156.9 ± 12.6 | 67.00 ± 22.00 | 14 |
| **PP Low** | 15 | 9 | 6 | 19.86 ± 0.94 | 72.97 ± 1.80 | 25.50 ± 1.05 | 165.2 ± 12.7 | 6.39 ± 0.60 | 9 |
| **PP High** | 15 | 7 | 8 | 18.49 ± 1.17 | 72.96 ± 2.83 | 24.62 ± 1.58 | 152.8 ± 12.9 | 66.40 ± 18.88 | 15 |

### 3.1. Blood lead levels (BLL)

Birds sampled at sites with high soil lead concentrations (mine or smelter vicinities) showed markedly elevated BLL. In the Broken Hill exposed group, BLL values ranged from 12.6 to 111.0 µg/dL (mean ± SD: 49.0 ± 31.4 µg/dL, n = 31). In the Port Pirie exposed group, blood lead ranged from 28.7 to 111.0 µg/dL (mean ± SD: 56.0 ± 22.9 µg/dL µg/dL, n = 25). BLL did not differ significantly between the two high-BLL groups (Welch t-test: p = 0.3). In the Broken Hill low-exposure group, lead was measured in 12 birds, and all birds returned values below the instrument’s lower detection limit (<5.97 µg/dL ICP-MS equivalent). In Port Pirie, blood lead was quantifiable in 9 of 12 low-exposure individuals, ranging from 5.97 to 40.1 µg/dL (mean ± SD: 10.9 ± 10.1). These results show that the low-exposure site classification accurately reflects minimal systemic lead uptake, particularly in Broken Hill, where contamination is more geographically constrained. In PP Low, BLLs were slightly higher than in BH Low but significantly lower than in PP High.

### 3.2. Blood haemoglobin decreases with increasing blood lead levels

Lead toxicity is well known to affect heme biosynthesis by inhibiting δ-aminolevulinic acid dehydratase (δ-ALAD), an essential enzyme in heme synthesis [44]. Blood haemoglobin (Hb) levels (g/L) were measured immediately after sampling using a HemoCue Hb 201+ instrument, and summary statistics of heme levels in all groups are provided in Supplementary Table S3. To assess the relationship between blood lead and haemoglobin in house sparrows from Broken Hill and Port Pirie, we employed quantile regression at the conditional median (τ = 0.5) rather than ordinary least squares, as residuals from the Broken Hill model violated the normality assumption (Shapiro-Wilk test). In Broken Hill, the relationship between blood lead and haemoglobin was significant: moving from the low-exposure median (5.97 µg/dL) to the high-exposure median (59.51 µg/dL) was associated with a predicted haemoglobin decrease of 11.0 g/L (β = −4.79 g/L per unit ln(blood lead), τ = 0.5, p = 0.0006) (Figure 3). In Port Pirie, the equivalent regression was not significant (β = −4.24 g/L per unit ln(blood lead), τ = 0.5, p = 0.15). When both towns were modelled together, the overall regression remained significant (β = −4.72 g/L per unit ln(blood lead), τ = 0.5, p = 0.0009), and the location × blood lead interaction was not significant (p = 0.86), indicating that the slopes did not differ statistically between towns, likely due to insufficient power to detect a difference at the available sample sizes. It should be noted that, by design, this study targeted birds at the extremes of lead exposure, without intermediate blood lead values, which limits continuous dose-response inference and renders the analysis more closely akin to a categorical comparison between exposure groups.

**Figure 2:**
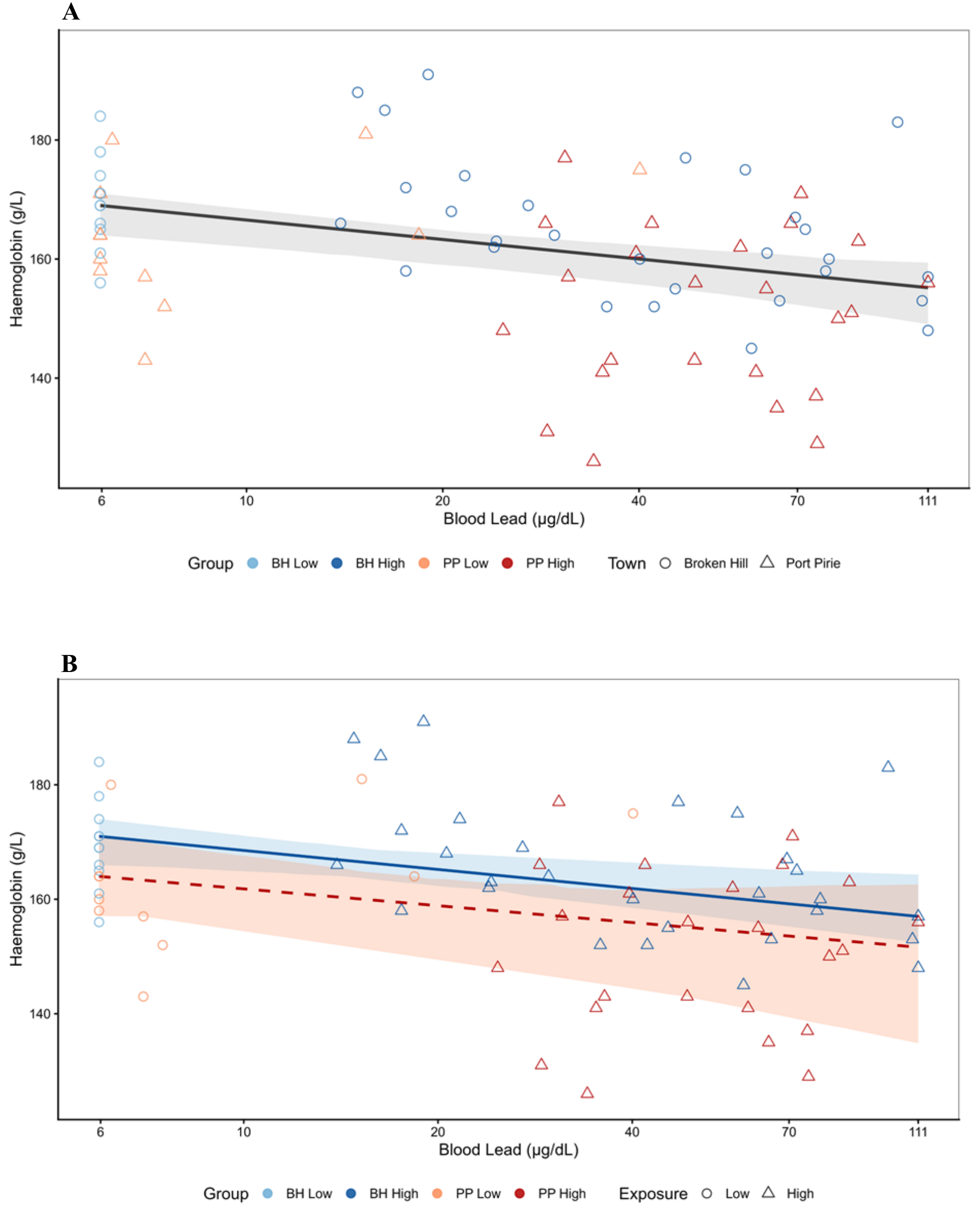
Relationship between blood lead concentration (µg/dL) and haemoglobin (g/L) estimated by quantile regression at the conditional median (τ = 0.5). Regression lines represent the conditional median, and shaded areas indicate 95% bootstrap confidence intervals. (a) Pooled model combining both towns (Broken Hill + Port Pirie); β = −4.72 g/L per unit ln(blood lead), p = 0.0009. (b) Town-stratified models; solid blue line = Broken Hill (β = −4.79 g/L per unit ln(blood lead), p = 0.0006), dashed red line = Port Pirie (β = −4.24 g/L per unit ln(blood lead), p = 0.149). Point colours indicate exposure group: dark blue = BH High, light blue = BH Low, dark red = PP High, light orange = PP Low. Blood lead values are displayed on a log-spaced axis in original µg/dL units following Gillings *et al.* correction. n = 54 adults (BH = 26, PP = 22; one Broken Hill High outlier excluded).

**Figure 3:**
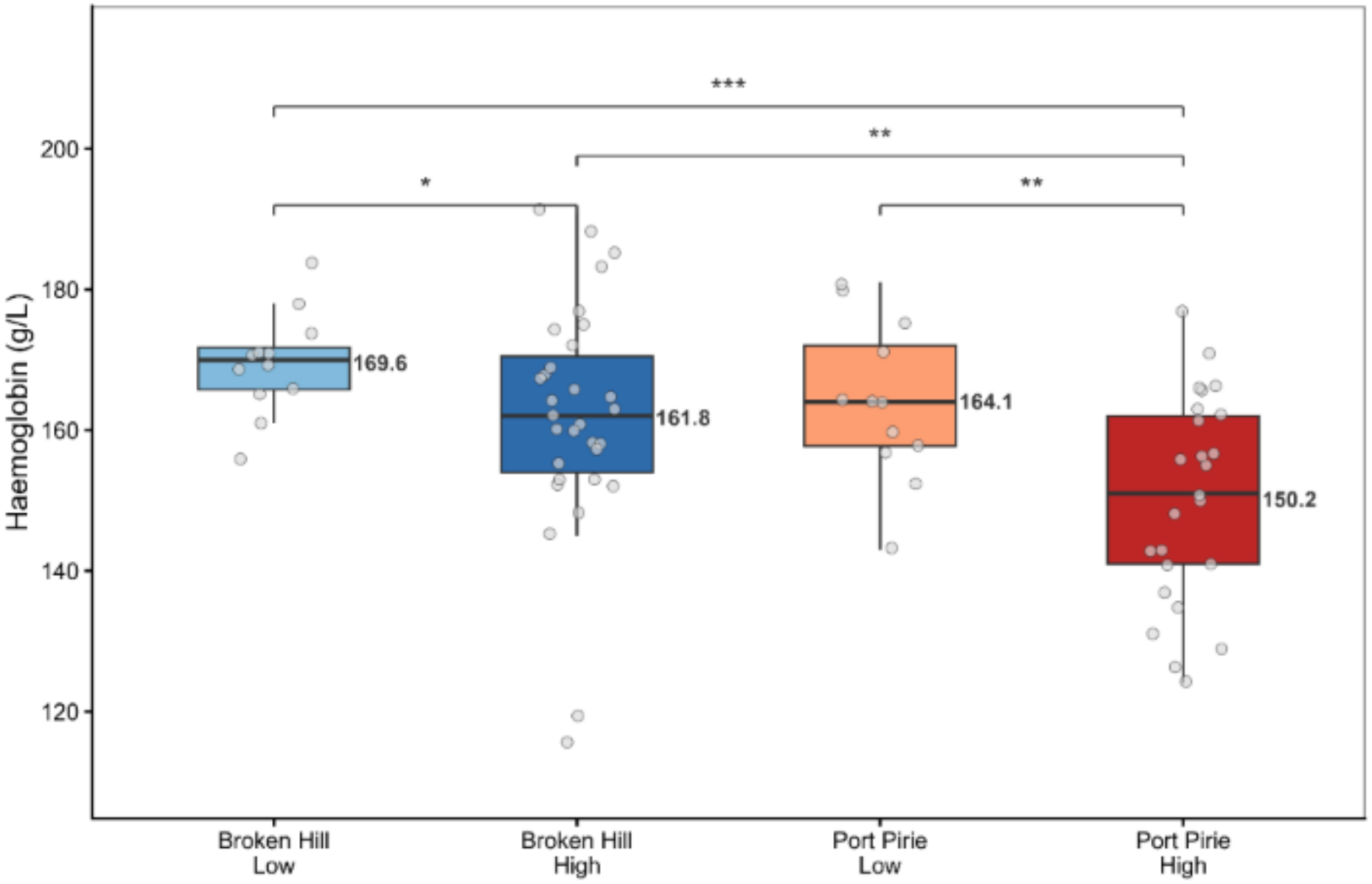
Haemoglobin concentrations (g/L) across exposure groups in Broken Hill and Port Pirie house sparrows. Boxes show the interquartile range, horizontal lines indicate the median, whiskers extend to 1.5 × IQR, and individual points represent single birds. Pairwise Welch t-tests (two-sided) were performed between all groups; only significant comparisons are shown (** p<0.01, *** p<0.001). BH Low (n=12, mean=169.6 g/L), BH High (n=31, mean=161.8 g/L), PP Low (n=12, mean=164.1 g/L), PP High (n=25, mean=150.2 g/L). BH = Broken Hill; PP = Port Pirie.

Across all lead exposure levels, Broken Hill birds maintained significantly higher haemoglobin concentrations than Port Pirie birds (BH mean = 164 g/L, n = 43 vs PP mean = 155 g/L, n = 37; Welch t-test: p = 0.007), which indicates a systemic between-site difference in haematopoietic capacity independent of lead exposure classification. Within Broken Hill, haemoglobin was significantly higher in low-exposure birds than high-exposure birds (mean 170 vs 162 g/L; Welch t-test: p = 0.04). Similarly, within Port Pirie, low-exposure birds had significantly higher haemoglobin than high-exposure birds (mean 164 vs 150 g/L; p = 0.004). Notably, BH HL birds (mean 162 g/L) maintained significantly higher haemoglobin than PP HL birds (mean 150 g/L; Welch t-test: p = 0.008), despite equivalent blood lead concentrations between the two high-exposure groups (Welch t-test: p = 0.3). This suggests that birds in Port Pirie experience an additional haemoglobin deficit. In contrast, BH HL and PP LL birds did not differ significantly in haemoglobin (Welch t-test: p = 0.6), indicating that the haemoglobin suppression associated with high lead exposure in Broken Hill is of comparable magnitude to the baseline haemoglobin deficit carried by minimally lead-exposed Port Pirie birds. This suggests that, despite suppression of heme synthesis, Broken Hill birds maintain higher heme levels than the Port Pirie birds.

### 3.3. Blood Proteomics data overview

To investigate proteome changes induced by lead toxicity in two isolated populations of house sparrows, we performed quantitative proteomic analysis using label-free data-independent acquisition (DIA) mass spectrometry. For proteomic analysis, we selected 15 blood samples from each group, prepared them to use an optimised dried blood spot method with the Mitra VAMS technique, and analysed them on an Exploris 480 mass spectrometer. An empirical spectral library was constructed using gas-phase fractionation (GPF) of pooled tryptic peptides. Library generation identified 3,480 proteins and was subsequently used as a spectral reference for protein identification in single-injection analyses.

Raw data quality control identified three samples with no recorded values, which were excluded from further analysis. These three samples belonged to the Port Pirie Low exposure group and likely failed during the desalting step. Reducing the remaining groups to n = 12 to equalise sample sizes was not pursued, as this would discard valid data and reduce statistical power without methodological justification; the differential abundance analysis framework used in this study (MSDAP) accommodates unequal group sizes by design through its underlying linear modelling and empirical Bayes moderation, and variance estimation is improved by retaining all available observations. Of the remaining 57 single-injection samples, 1,955 protein groups were identified, of which 848 (43.4%) were identified without missing values across all samples. To assess quantification reproducibility, six quality control samples were analysed in parallel, yielding a median coefficient of variation (CV) of 16.3% at the protein group level (Figure 4A). Our VAMS-based blood collection approach incorporates a depletion step to remove highly abundant proteins, most notably haemoglobin, ensuring consistent removal across all samples and minimising bias that could arise from differential washing efficiency. Haemoglobin abundance was consistent in all groups (Figure 4B). Importantly, while circulating haemoglobin concentrations differed between exposure groups (as measured by HemaCue), the LiCl washing protocol depleted haemoglobin to uniformly low levels across all samples regardless of starting concentration, ensuring that biological differences in haemoglobin abundance did not introduce quantification bias into the proteomic dataset.

**Figure 4:**
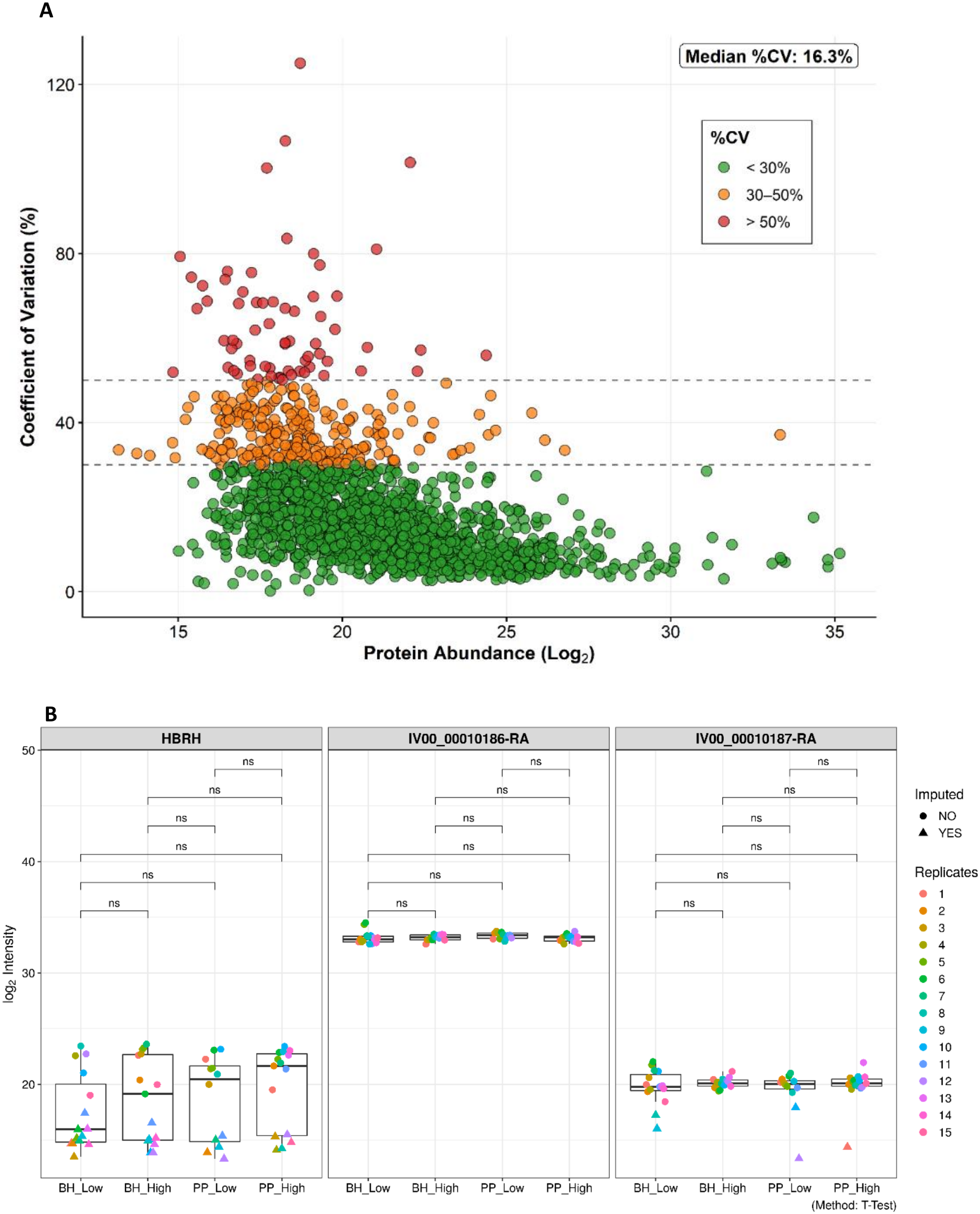
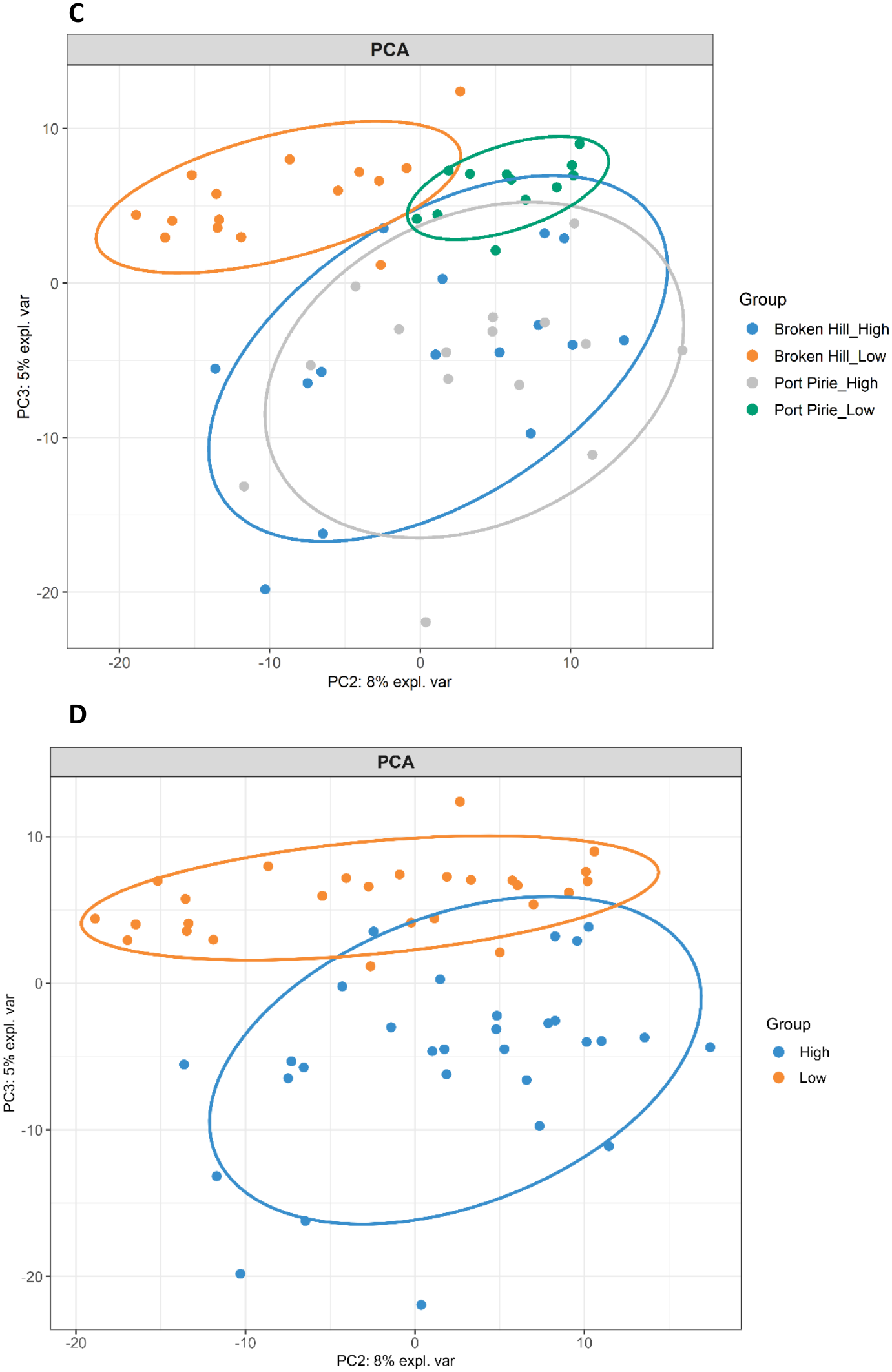
Overview of blood proteomics data quality and sample structure. A) Protein abundance versus percentage coefficient of variation (%CV) across all quantified protein groups. The median %CV was 16.3%, with most proteins falling below the 30% threshold. B) Haemoglobin proteins were uniformly washed. Three subunits of haemoglobin indicate that the haemoglobin was washed uniformly. Significance is based on t-test. C) Unsupervised principal component analysis (PCA) of the 1,000 most variable protein features. Class separation was observed on PC2 and PC3, with low-exposure birds from Broken Hill and Port Pirie showing partial separation, reflecting site-specific differences in baseline proteome composition. High-exposure birds from both towns clustered together without apparent separation. D) PCA Proteomics data overview extended. The high lead and low lead separate on component 3.

Unsupervised principal component analysis (PCA) was performed using the 1,000 most variable features. Class separation was observed on PC2 and PC3, with clear segregation of low-and high-exposure birds along these components (Figure 2C and 2D). Within the low-exposure group, birds from Broken Hill and Port Pirie showed partial separation, suggesting site-specific differences in baseline proteome composition under minimal lead burden. In contrast, high-exposure birds from both towns clustered together without apparent separation.

### 3.4. Proteomic responses to lead exposure converge on RNA processing

Differential abundance analysis of Broken Hill birds revealed 91 differentially abundant proteins, of which 28 were significantly higher abundant, and 63 were significantly lower abundant in the high blood lead group (Figure 5A and Supplementary Table S4). Among the differentially abundant proteins, several canonical oxidative stress response proteins were identified. Peroxiredoxin 6 (PRDX6), BTB and CNC homology 1 (BATCH1), and Sequestosome 1 (SQSTM1, also called p62) were elevated in the high lead-exposure group, whereas Peroxiredoxin 4 (PRDX4) was decreased, consistent with oxidative inactivation and consumption of the peroxiredoxin antioxidant defence under conditions of chronic lead-induced oxidative stress [45, 46]. In the Broken Hill high lead exposure group, multiple subunits of the TRiC/CCT chaperonin complex (TCP1, CCT2, CCT3, CCT5, CCT6, CCT7, CCT8) were among the most consistently increased abundance proteins. CCT proteins have been independently identified as hub proteins in a cadmium-exposed human cohort, indicating that upregulation of this pathway is a conserved molecular hallmark of heavy-metal proteotoxicity across species and metals [47].

**Figure 5:**
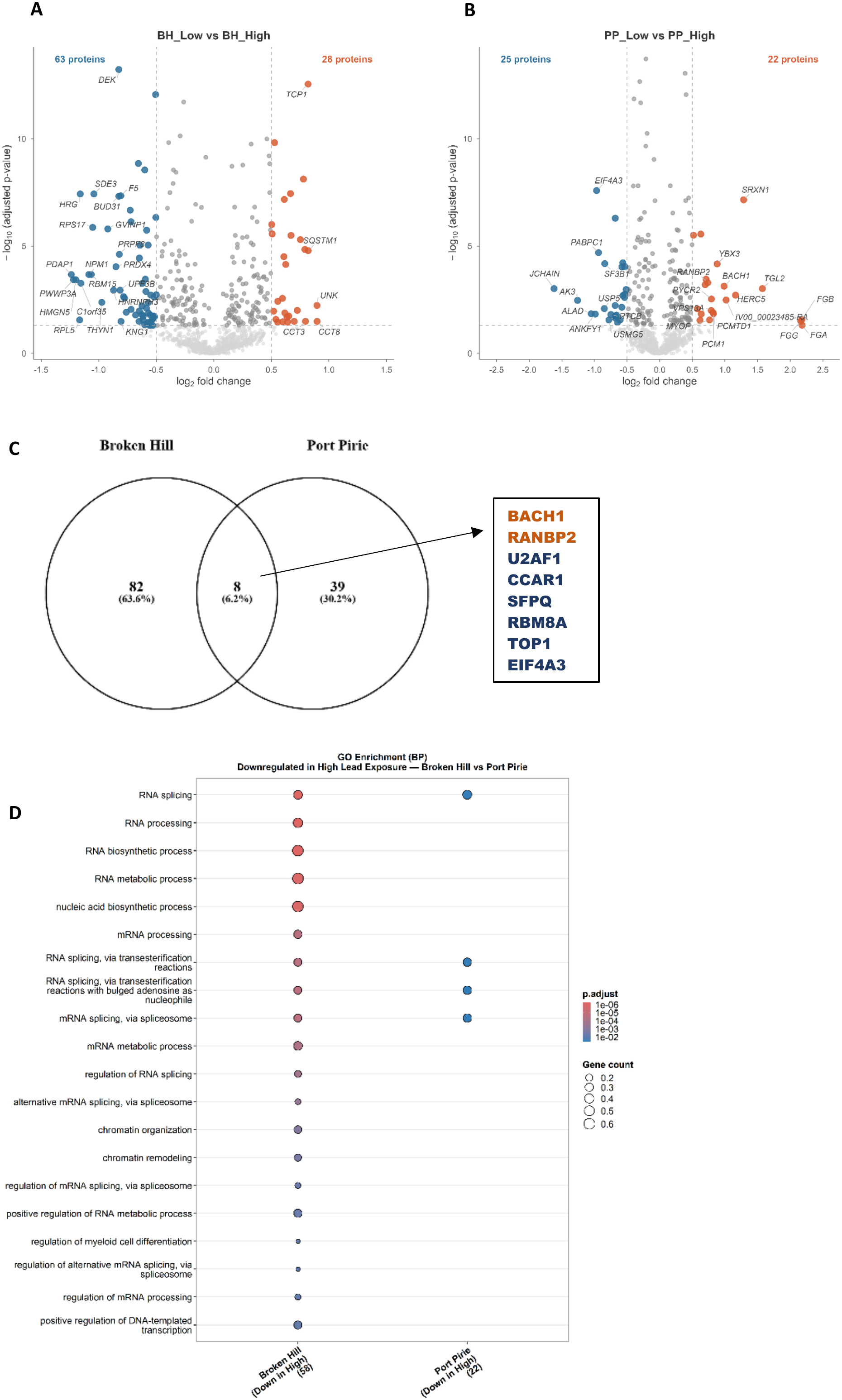
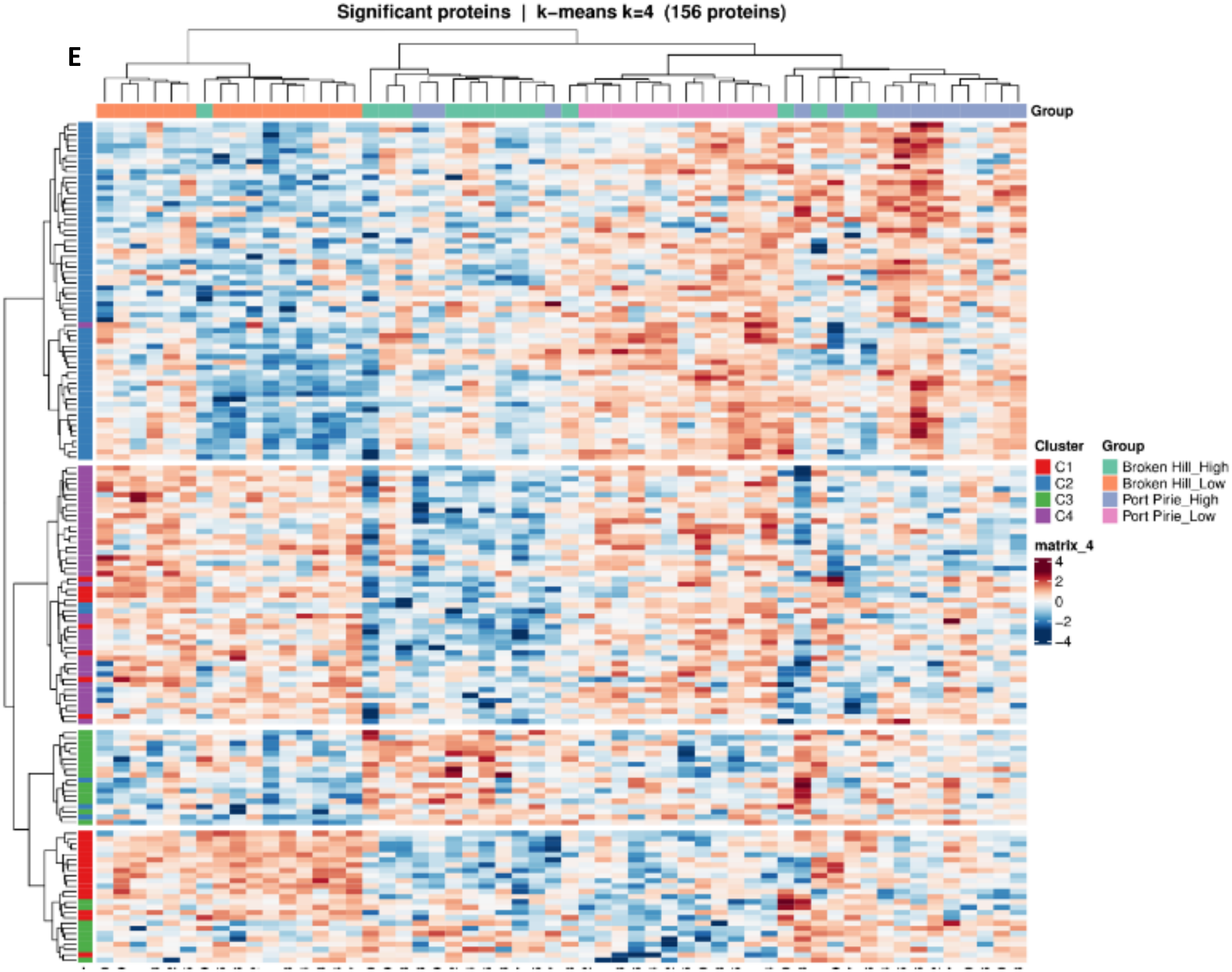
Differential protein abundance between low- and high-exposure groups within each town. Volcano plots display log₂ fold change on the x-axis and −log₁₀ adjusted p-value on the y-axis; dashed lines indicate significance thresholds. A) Volcano plot for Broken Hill (High vs Low): proteins with significantly higher abundance in the high-exposure group are shown in red and proteins with significantly lower abundance are shown in blue. B) Volcano plot for Port Pirie (High vs Low), using the same colour scheme. C) Venn diagram showing the overlap of significantly differentially abundant proteins between the two towns. Red segments represent proteins with higher abundance in high-exposure birds, and blue segments represent proteins with lower abundance in high-exposure birds. D) Gene ontology enrichment of Biological Processes for lower abundant proteins in both towns. E) Heatmap of significantly differentially abundant proteins identified using one-way ANOVA. Rows represent individual proteins and columns represent samples, both ordered by hierarchical clustering. Protein abundance values are displayed as z-scores (row-scaled) to enable cross-protein comparison of relative abundance patterns. Row annotations indicate protein clusters (C1–C4) identified by k-means clustering.

In Port Pirie sparrows, 47 proteins were differentially abundant, with 22 elevated in abundance and 25 reduced in birds with high blood lead levels (Figure 5B and Supplementary Table S5). Among the differentially abundant proteins, several canonical oxidative stress response proteins were identified. SRXN1 (Sulfiredoxin-1) and TXN2 (Thioredoxin 2) were elevated in the high lead-exposure group, reflecting an active antioxidant rescue response, whereas GPX1 was notably decreased, consistent with oxidative inactivation and glutathione peroxidase defence depletion under conditions of chronic lead-induced oxidative stress [48, 49]. The elevation of SRXN1, which specifically repairs hyperoxidised peroxiredoxins, further supports the interpretation that the peroxiredoxin antioxidant system is being overwhelmed and consumed, corroborating the PRDX4 depletion observed in Broken Hill birds [50]. In the Port Pirie high group, PRDX4 showed a statistically significant decrease (q-value < 0.05). However, the magnitude of change did not meet the fold-change threshold (log2FC = 0.40) and was therefore not classified as differentially abundant. Notably, δ-aminolevulinic acid dehydratase (δ-ALAD), the direct enzymatic target of lead toxicity, was significantly reduced in Port Pirie birds but showed no significant change in Broken Hill birds, suggesting differential sensitivity of this canonical lead-response protein between the two populations. Eight proteins were shared between the two comparisons; two of which had elevated abundance levels in high lead groups, and six were less abundant in the high blood lead groups in both towns (Figure 5C). The direction of change was consistent across both towns for all eight shared proteins, suggesting that their expression responses are independent of geographic location and reflect a common molecular response to elevated blood lead levels. The two shared proteins increased in abundance were BACH1 and RANBP2 (Ran-binding protein 2, also called Nup358), while the six shared proteins decreased in abundance were U2AF1, CCAR1, SFPQ, RBM8A, TOP1, and EIF4A3.

Gene enrichment analysis for biological processes was performed on the differentially abundant proteins (Supplementary Tables S6 to S9). Among proteins increased in abundance in Broken Hill sparrows, the most significantly enriched biological processes included regulation of telomere maintenance and sperm binding to zona pellucida. Both terms are driven by the same set of genes, i.e., the TRiC/CCT complex proteins. The complex plays an important role in protein quality and is responsible for the correct folding of approximately 10% of all cytosolic proteins [51]. The complex also plays a role in regulating telomere maintenance by folding TCAB1 [52]. We have previously reported severe telomere disruption in Broken Hill house sparrows exposed to lead toxicity [53], and activation of this complex may play a role in protein quality control and telomere elongation. In contrast, proteins increased in abundance in Port Pirie sparrows were enriched for distinct biological processes, including blood coagulation and related haematological processes. There was no overlap with those in Broken Hill birds, suggesting a town-specific proteomic response to lead at the level of upregulation. At the level of downregulation, however, the two towns showed a more convergent response. In both Broken Hill and Port Pirie sparrows, proteins decreased in abundance were enriched for RNA processing and splicing, suggesting that impaired post-transcriptional regulation is a shared molecular consequence of lead exposure (Figure 5D). Because erythrocytes make up the vast majority of blood cells, and avian erythrocytes, although nucleated, have reduced transcriptional and translational activity compared with other cell types, this decrease may reflect a shift in the age structure of the circulating erythrocyte population rather than a direct effect on RNA processing. For example, lead-induced haemolysis and impaired erythropoiesis could alter the proportion of mature to newly produced cells. Whether the loss of RNA processing proteins reflects altered erythrocyte turnover or direct lead toxicity remains to be determined.

To further characterise proteomic changes across all four groups, one-way ANOVA with Tukey post hoc testing and Benjamini-Hochberg correction was performed, identifying 156 proteins with significantly different abundances across comparisons (BH adjusted p < 0.05). Hierarchical clustering of these proteins and samples, visualised as a heatmap, produced a clear separation of the four groups with only minor misclassification between the two high-exposure groups across towns (Figure 5E). Cluster 2 (C2) appeared to reflect a town-level difference in protein abundance that was largely independent of lead exposure status, while no significant biological process was enriched. Cluster 4 (C4), however, displayed a coordinated response to lead exposure, with high-exposure groups from both towns showing consistently lower abundance relative to their respective low-exposure counterparts. GO enrichment of C4 proteins revealed significant enrichment in RNA splicing, spliceosome-mediated mRNA splicing, and mRNA processing (Supplementary Table S10). Seven proteins were identified with significantly different abundances across groups (BH-corrected p < 0.05) that were shared between the two towns: AFG3L2, LMNB2, RPS27A, CYB5R3, PRDX4, ZNF326, and SNRPD1. Although not all proteins met the fold-change threshold in each comparison, the direction of change was consistent across towns. Of these, AFG3L2, a mitochondrial quality-control protease, was the only protein moderately elevated in response to lead exposure, whereas the remaining six were uniformly decreased in abundance. When these seven proteins were combined with the eight previously identified through the MS-DAP multi-model analysis, yielding a combined set of 12 unique proteins, GO enrichment analysis revealed significant enrichment in RNA splicing (FDR = 0.00061).

### 3.5. Divergent proteomic response to lead between Broken Hill and Port Pirie

To identify proteins exhibiting divergent lead responses between populations, a protein-wise linear model including Location, Lead level, and their interaction (Location × Lead level) was fitted. Detection of interaction effects is often underpowered in high-dimensional proteomic datasets with moderate sample sizes (n = 12–15 per group); therefore, empirical Bayes moderation was applied to borrow information across proteins, resulting in more stable variance estimates and improved statistical sensitivity. This approach substantially increased the number of nominally significant interactions, though many proteins still did not survive FDR correction. Results are therefore reported at nominal p-value thresholds (p < 0.05), and the identified candidates are interpreted as hypothesis-generating rather than formally confirmed.

The interaction analysis identified 137 proteins whose lead response differed significantly between BH and PP (p < 0.05, |log2FC| > 0.25), which are shown as a scatterplot in Figure 6A, with a full list provided in Supplementary Table S11. At the pathway level, the 26S proteasome regulatory particle showed a coordinated divergence. The AAA-ATPase ring subunits (PSMC1, PSMC2, PSMC3, PSMC5, PSMC6) were consistently increased in abundance in BH high lead exposure areas and were unchanged or suppressed in PP. Three proteasome regulatory subunits showed individually significant divergent responses (PSMC5 p=0.008, PSMC3 p=0.029, PSMD2 p=0.042), with BH birds indicating increased abundance and PP birds showing no change or mild suppression. Three additional subunits (PSMC1, PSMC2, PSMC6) showed the same directional pattern but did not reach nominal significance (p=0.07 to 0.09). The concordant directionality across all six subunits is notable. The stress-inducible HSP70 family members HSPA2, HSPA4 and HSPA8 were all increased in abundance in BH lead-exposed birds and suppressed in PP. The TRiC/CCT complex subunit CCT5 showed the same pattern. Heat shock proteins act as molecular chaperones that assist in the correct folding of proteins under challenging conditions [54]. Lead is known to disrupt protein structural integrity through direct binding to thiol groups and displacement of essential metal cofactors, and the increased abundance of these chaperones in BH birds may help maintain proteome integrity by refolding damaged proteins before they aggregate [55]. Heat shock proteins have also been recognised as evolutionary capacitors, buffering the phenotypic effects of genetic variation and potentially facilitating adaptation to novel environments [56, 57].

**Figure 6:**
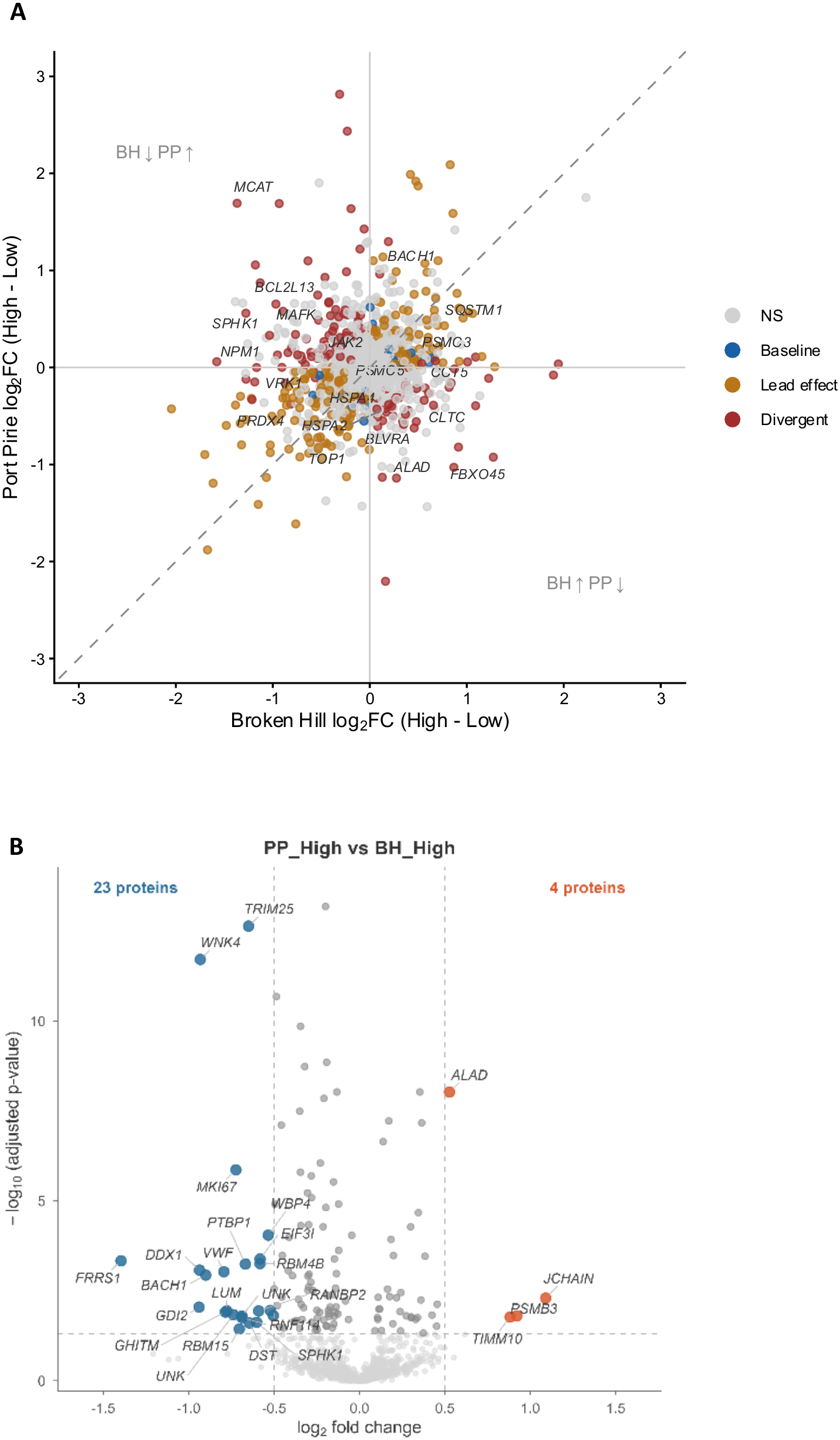
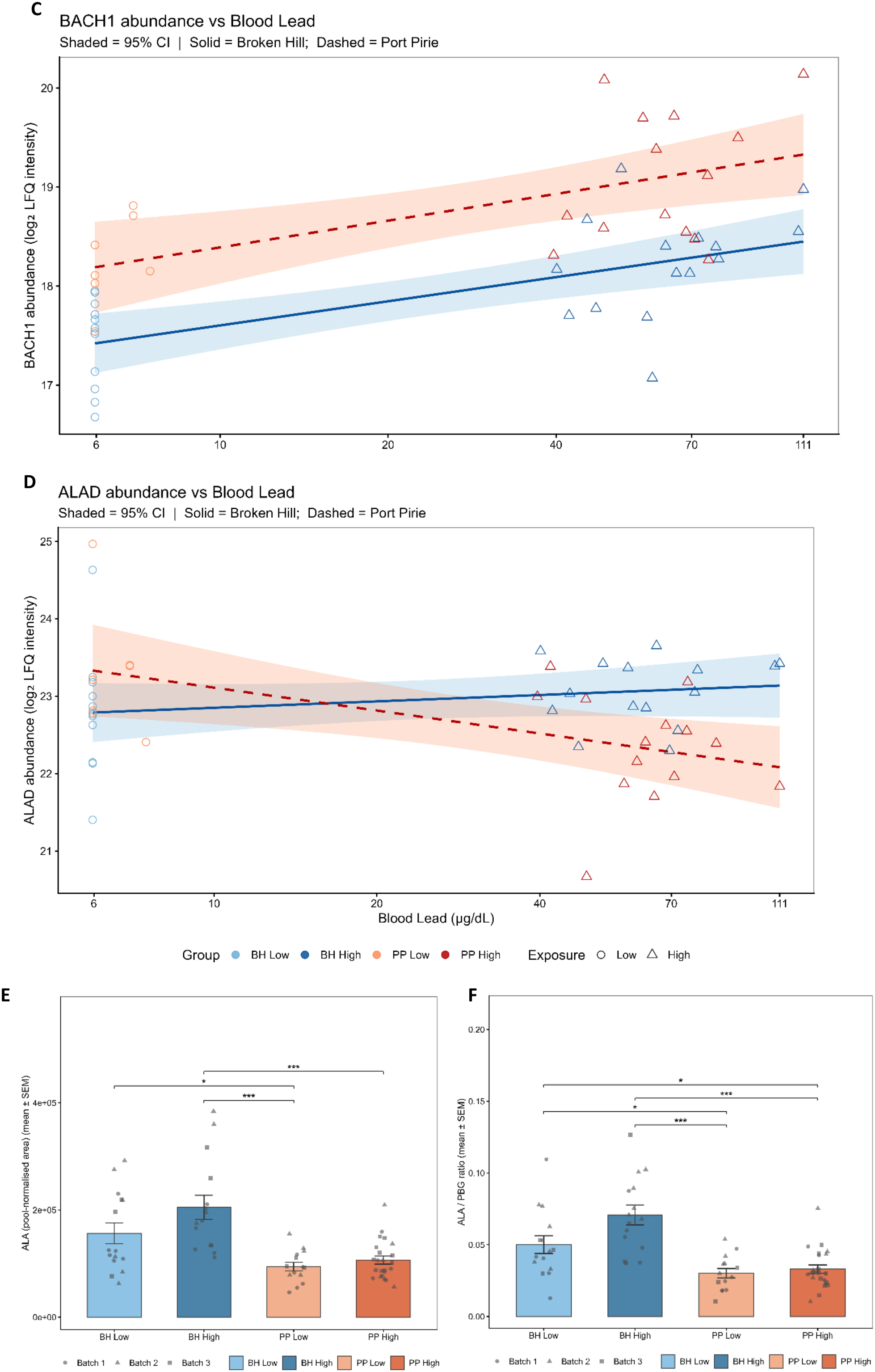
Proteomic response to lead exposure in high-exposure birds from Broken Hill and Port Pirie. A) Scatterplot of protein-level lead response in Broken Hill versus Port Pirie. Each point represents a single protein, with the x-axis displaying the log₂ fold change between high and low exposure groups within Broken Hill and the y-axis displaying the equivalent comparison within Port Pirie. Proteins are coloured by response category: baseline (no significant change in either town), lead effect (significant change in the same direction in both towns), and divergence (significant change in opposing directions between towns). Proteins falling along the diagonal respond similarly to lead in both populations, whereas proteins deviating from the diagonal indicate population-specific responses. B) Volcano plot comparing protein abundance between Broken Hill High and Port Pirie High exposure groups. Proteins with significantly higher abundance in Broken Hill High birds are shown in red, and proteins with significantly higher abundance in Port Pirie High birds are shown in blue. C) Linear regression of blood lead concentration against BACH1 (BTB and CNC homology 1) protein abundance (log2 scaled). The solid blue line represents Broken Hill, and the dashed red line represents Port Pirie; shaded areas indicate 95% confidence intervals. D) Linear regression of blood lead concentration against ALAD (δ-aminolevulinic acid dehydratase) protein abundance, using the same colour scheme. E) ALA levels in blood, ALA was measured as the mono-NPH derivative (precursor m/z 265.10, quantifier transition 265.10 to 152.15). Significance brackets show Tukey HSD adjusted p-values from a two-way ANOVA on log₂-transformed values (Location × Lead level, Type II SS). F) ALA/PBG ratio in whole blood as a functional readout of ALAD enzyme inhibition. The ratio was calculated from pool-normalised peak areas of 5-aminolevulinic acid (ALA, quantifier 265.10 to 152.15) and porphobilinogen (PBG, quantifier 359.20 to 122.05) for each individual bird. A higher ratio indicates greater accumulation of the ALAD substrate (ALA) relative to its product (PBG). Significance brackets show Tukey HSD adjusted p-values from a two-way ANOVA on log₂-transformed ratios (Location × Lead level, Type II SS).

At the protein level, the E3 ubiquitin ligase FBXO45 exhibited one of the largest interaction effects in the dataset, increasing markedly in BH birds (+1.27 log2FC) while decreasing in PP birds (−0.93 log2FC). However, the most biologically informative interaction effects were observed among proteins associated with haem metabolism. ALAD (δ-aminolevulinic acid dehydratase), the direct molecular target of lead toxicity, showed a substantial decrease in abundance in lead-exposed PP birds (−1.13 log2FC) but remained largely unchanged in BH birds (+0.13 log2FC). BLVRA (biliverdin reductase A), which catalyses the final step of haem degradation, was likewise reduced in PP birds exposed to lead (−0.55 log2FC) but showed no comparable response in BH birds, indicating a population-specific disruption of haem homeostasis. To investigate the downstream consequences of this ALAD divergence, we directly compared the two high-lead groups (PP High vs BH High) and identified 4 proteins with significantly higher abundance in BH high lead birds (TIMM10, PSMB3, JCHAIN and ALAD), and 23 proteins with higher abundance in PP high lead birds (Figure 6B). Among the proteins elevated in PP birds, BACH1 was of biological interest given its mechanistic link to ALAD through haem synthesis [58].

ALAD is one of the most sensitive enzymatic targets of lead toxicity. Lead directly inhibits ALAD activity, causing accumulation of its substrate aminolevulinic acid (ALA) and reducing haem biosynthesis. Free intracellular haem acts as a negative regulator of BACH1 by binding to it and triggering its proteasomal degradation [57]. BACH1, in turn, forms a heterodimer with MAFK that binds to antioxidant response elements (AREs) and Maf recognition elements (MAREs), repressing the transcription of haem-responsive genes, including globins and HO-1. When lead inhibits ALAD and depletes haem, BACH1 is no longer efficiently degraded and accumulates within the cell, reducing the production of globin chains available for haemoglobin assembly [59, 60]. The consequence is a dual impairment of haemoglobin synthesis: reduced haem supply from ALAD inhibition and reduced globin availability from BACH1-mediated transcriptional repression, both arising from the same initiating event. Lead essentially hijacks the cell’s own haem-sensing machinery, converting a physiological regulatory feedback loop into a pathological cascade that amplifies the haematological consequences of its own toxicity.

Consistent with this model, the pairwise comparison confirmed that both BACH1 and its binding partner MAFK were significantly more abundant in PP High relative to BH High birds. We performed linear modelling with BLLs and BACH1 abundance was significantly positively associated with blood lead levels in both Broken Hill (R² = 0.40, p = 0.0002) and Port Pirie (R² = 0.40, p = 0.0028), consistent with progressive BACH1 stabilisation as lead-induced haem depletion worsens with increasing exposure (Figure 6C). The town by lead interaction for BACH1 was non-significant (p = 0.8). However, BACH1 abundance remained substantially lower across all lead levels in BH birds than in PP birds, suggesting that haem depletion and consequent globin suppression are more severe in PP birds. For ALAD, the model revealed a significant negative relationship with blood lead in PP birds (R² = 0.30, p = 0.009) but no significant relationship in BH birds (R² = 0.05, p = 0.255), with a significant town by lead interaction (p = 0.0031) (Figure 6D).

Together, these results support a model in which lead-induced inhibition of ALAD affects multiple downstream components of erythroid physiology through a common regulatory pathway. Reduced ALAD activity would be expected to decrease haem synthesis, stabilise BACH1, and promote formation of the BACH1–MAFK repressor complex, thereby limiting globin gene expression and haemoglobin production. Elevated BACH1 activity may additionally suppress antioxidant response pathways, potentially increase oxidative stress and alter protein quality-control mechanisms. In BH birds, maintenance of ALAD abundance may preserve haem synthesis and haem-dependent BACH1 turnover, reducing activation of these downstream responses.

### 3.6. Metabolite profiling reveals functional ALAD inhibition despite maintained protein abundance

To determine whether the observed divergence in ALAD protein abundance reflected functional differences in enzyme activity, 5-aminolevulinic acid (ALA) and porphobilinogen (PBG) concentrations were measured in whole blood as direct markers of ALAD substrate accumulation and product formation *in vivo*. We reasoned that if ALAD protein levels are comparable between towns, yet the enzyme is catalytically inhibited by lead, ALA should accumulate upstream while PBG, the immediate product of ALAD, should remain unchanged or decline. An active enzyme would be expected to convert ALA to PBG efficiently, yielding a low ALA/PBG ratio, whereas inhibition would elevate the ALA/PBG ratio.

Blood ALA levels were significantly higher in Broken Hill birds regardless of lead exposure level (Location: p < 0.001), with both BH Low and BH High groups exceeding their Port Pirie counterparts (Figure 6E). PBG concentrations showed no significant differences between any groups (Location: p = 0.49, Lead: p = 0.37). The ALA/PBG ratio confirmed this pattern, with Broken Hill birds retaining substantially higher ALA relative to PBG than Port Pirie birds (Location: p < 0.001, Lead: p = 0.021; Figure 6F). Notably, even the low-exposure Broken Hill group showed a significantly elevated ALA/PBG ratio compared to Port Pirie controls (1.6-fold, p = 0.025), indicating that ALA accumulation in Broken Hill birds is not solely a consequence of acute lead exposure but may reflect a baseline difference in ALAD functional capacity or ALA clearance between the two populations.

These findings are particularly interesting because ALA is itself a pro-oxidant molecule. At elevated concentrations, ALA undergoes auto-oxidation, generating reactive oxygen species that exacerbate oxidative damage independently of the direct effects of lead. The persistent elevation of blood ALA in Broken Hill birds, even at low exposure levels, suggests that these birds experience chronic oxidative stress from ALA accumulation in addition to the direct toxicity of lead. The elevated ALA may also drive the observed upregulation of ALAD protein in Broken Hill blood as a compensatory response; whereby increased enzyme abundance partially offsets catalytic inhibition to maintain flux through the haem biosynthesis pathway.

### 3.7. Lead changes mitochondrial activity in blood cells

BACH1 activation under lead toxicity is known to repress mitochondrial activity [61] and, in cancer cells, it promotes glycolysis while repressing the transcription of the tricarboxylic acid (TCA) cycle and the electron transport chain (ETC) [62, 63]. However, no significant changes in proteins associated with mitochondrial activity were identified in our blood proteomics dataset. To further investigate mitochondrial function, we performed a citrate synthase (CS) activity assay, which serves as an established proxy for mitochondrial volume [43]. CS activity did not differ significantly between high-and low-lead groups within either town individually, though Port Pirie approached significance (p = 0.075) with higher activity in lead-exposed birds (Figure 7A). When data were pooled across locations, the overall comparison between high-and low-lead groups showed a near-significant elevation of CS activity in lead-exposed birds (p = 0.052). This trend may suggest that lead exposure is associated with changed mitochondrial activity in sparrow blood, warranting further investigation.

**Figure 7.**
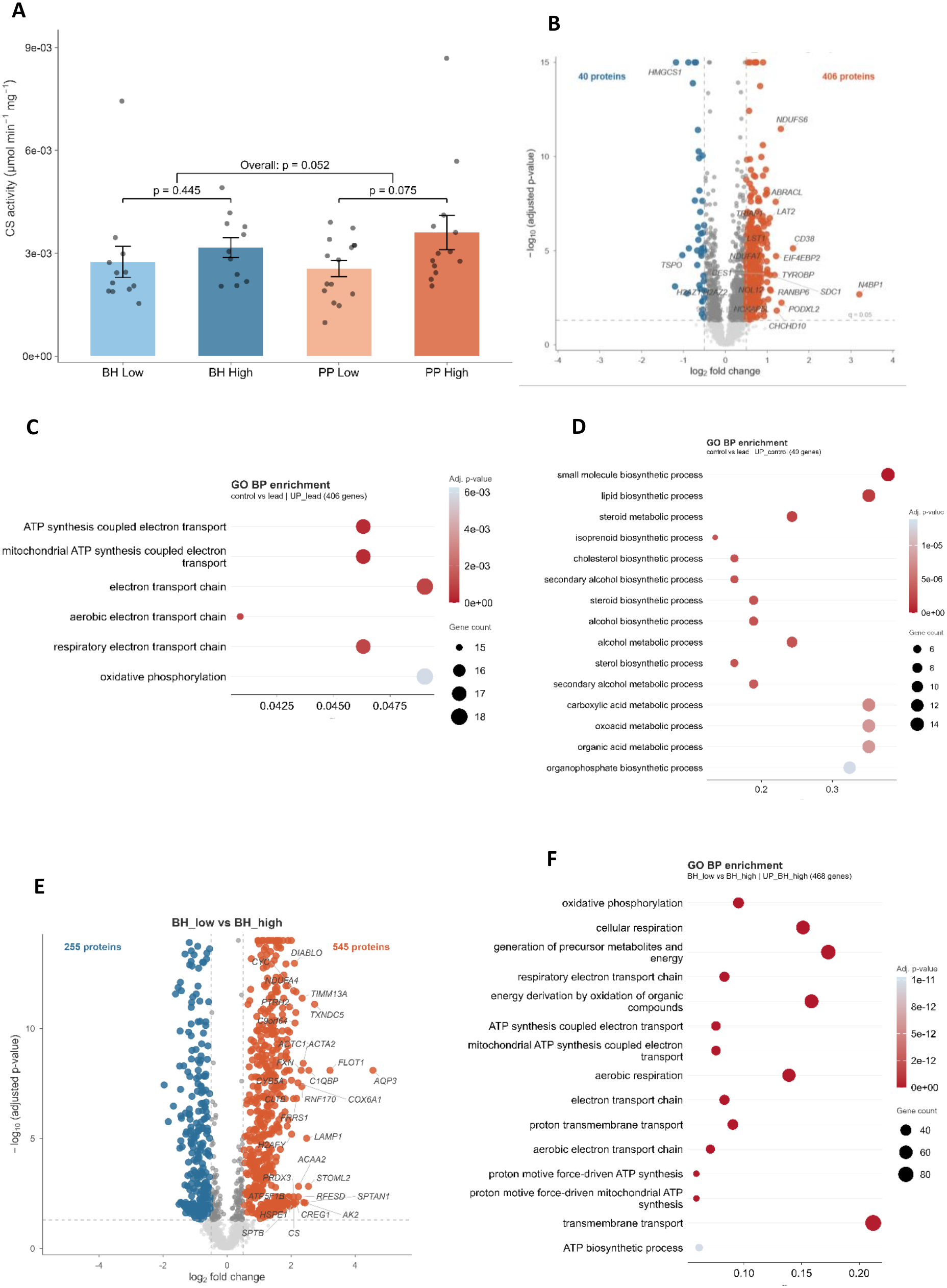
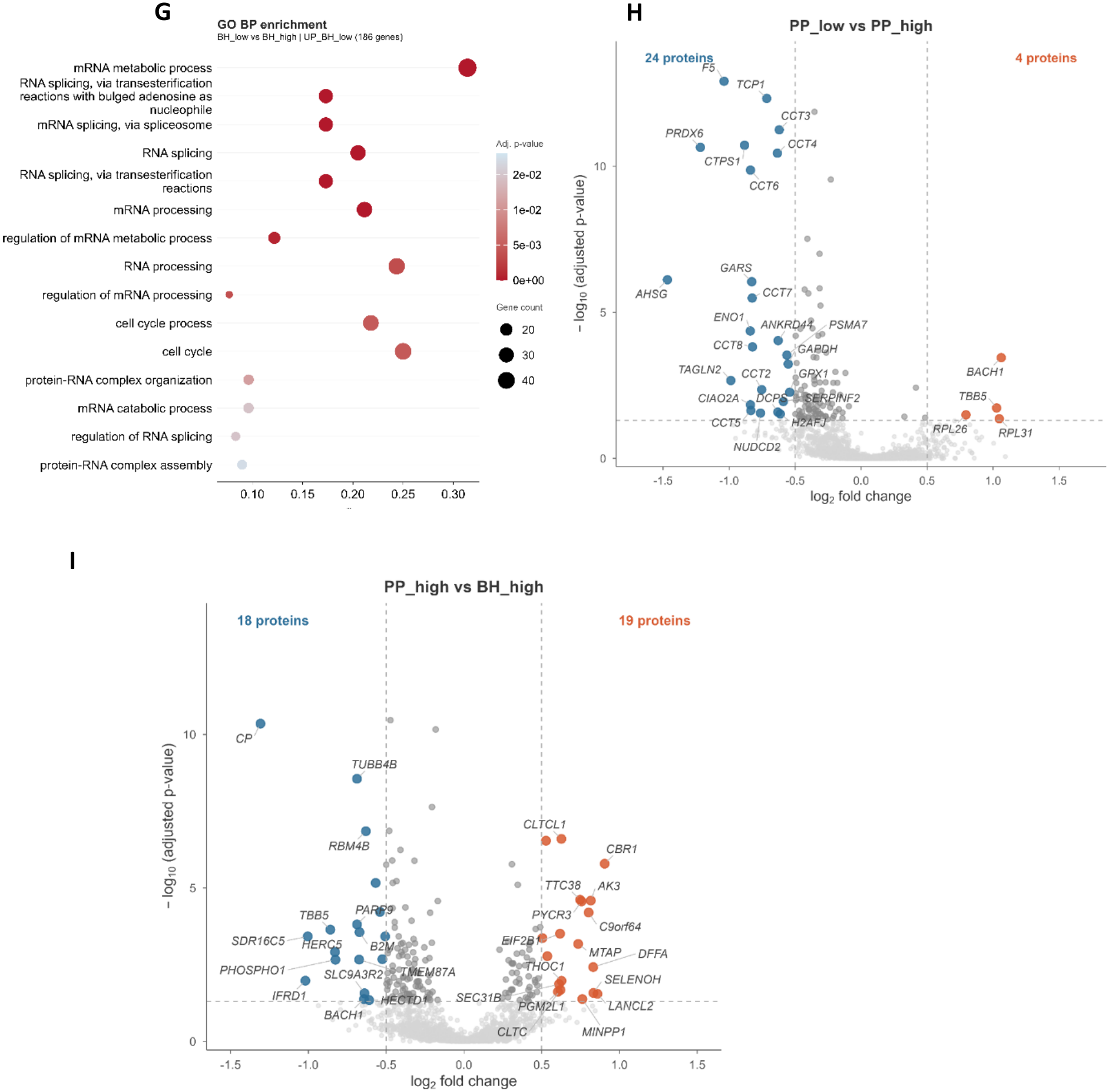
Mitochondrial activity in response to lead. A) Citrate synthase activity for samples with %CV<20 (n=51). Welch’s t-test was performed; error bars represent standard error of the mean (SEM), and p-values are printed. B) Volcano plot of differentially abundant proteins in the HL-60 cell line. C) Gene Ontology of Biological Processes for the elevated proteins in response to lead. D) Gene Ontology of Biological Processes for the decreased proteins in response to lead. E) Volcano plot of differentially abundant proteins identified from VAMS tip proteomics collected during the Broken Hill field trip, comparing high-versus low-exposure birds. Proteins with significantly higher abundance in the high-exposure group are shown in red and proteins with significantly lower abundance are shown in blue. F) Gene Ontology (GO) enrichment analysis of proteins with significantly higher abundance in Broken Hill high-exposure birds from secondary cohort. G) GO enrichment biological processes Broken Hill high vs Broken Hill low for proteins with higher abundance in low birds. H) Differential abundance analysis of Secondary Cohort for Port Pirie. I)Differential abundance analysis of Secondary Cohort for High lead birds from two towns.

### 3.8. In vitro lead exposure confirms direct mitochondrial impact independent of erythroid-specific responses

To investigate whether lead exposure directly impacts the mitochondrial proteome, we used the HL-60 promyelocytic cell line as an in vitro model. While a single cell line cannot fully represent the heterogeneous cell, populations present in whole blood, it provides a controlled system to isolate lead-specific proteomic changes from the environmental and genetic confounders inherent to field-collected samples. HL-60 cells were treated with 100 µg/dL lead acetate for 24 hours and analysed by DIA proteomics. Around 6,700 proteins were identified; of these, 5,000 passed quality filtering and were compared with untreated controls using the MSqRob model. A total of 446 proteins were significantly differentially abundant, with a striking asymmetry: 406 were more abundant in the lead-treated group and only 40 were decreased (Figure 7B). Gene ontology analysis of the proteins increased in abundance revealed that the top enriched biological processes were electron transport chain and oxidative phosphorylation (Figure 7C), confirming that lead directly impacts mitochondrial function and the oxidative stress response even in the absence of chronic exposure. Proteins that decreased under lead treatment were enriched in small-molecule biosynthesis, lipid metabolism, and sterol biosynthetic processes (Figure 7D), suggesting a metabolic trade-off in which biosynthetic capacity is sacrificed to support the mitochondrial stress response. ALAD and BACH1 were not significantly altered in HL-60 cells, which may be due to haem-BACH1 cascade being specific to erythroid lineage cells, or the duration of exposure to lead in birds.

### 3.9. Secondary cohort confirms key proteomic signatures under independent analytical conditions

An additional subset of field trip samples was prepared independently and analysed on an Orbitrap Astral mass spectrometer to assess whether key findings from the primary cohort were reproducible under different sample preparation and instrumentation conditions. This secondary cohort comprised 23 samples with an unbalanced design (BH Low, n = 4; BH High, n = 4; PP Low, n = 3; PP High, n = 12), reflecting opportunistic use of remaining material from our field trip. Two PP High samples were removed due to low protein identifications (cutoff: 2 SD below the median), leaving 21 samples for analysis. Lead levels were available only for the high exposure groups, and there was no significant difference between BH High (mean ± SD: 13.3 ± 3.15 µg/dL, n = 3) and PP High (mean ± SD: 17.3 ± 11.4 µg/dL, n = 10), p-value = 0.3377 (Welch t-test). We expected limited statistical power and fewer significant proteins due to lower lead exposure in this cohort compared to the original study. These samples were analysed on an Orbitrap Astral mass spectrometer, which identified 4239 proteins in total, compared with 1956 in our main cohort (a 2.17-fold increase). These gains are attributable to the superior sensitivity of the Orbitrap Astral instrumentation.

When we compared BH High versus BH Low sparrows in this dataset, we identified 800 significantly differentially abundant proteins (Figure 7E). Of these, 20 proteins overlapped with the primary BH cohort analysis, with 18 showing changes in the same direction. Gene ontology enrichment of proteins increased in abundance revealed significant enrichment for oxidative phosphorylation and ATP synthesis–coupled electron transport (Figure 7F), consistent with the CS activity assay and HL-60 cell line results. Decreased-abundance proteins were enriched in mRNA metabolic processes and RNA splicing, corroborating the primary dataset (Figure 7G). These findings are consistent with the primary cohort and provide additional support for the conclusion that mitochondrial activity is upregulated under lead stress and that RNA splicing is significantly altered by lead toxicity. Neither BACH1 nor ALAD showed significant changes in this BH comparison. When we performed the divergence analysis using this cohort, the results were very similar to those from the primary cohort, with chaperones (HSPA9, HSPA2, HSPA8, CCT2, CCT4, CCT5, CCT7, CCT8) and the proteasome (PSMA5, PSMA2, PSMD1, PSMB2, PSMC6) increased in BH birds as a result of exposure, compared with a decrease in PP birds. The divergence in these pathways in BH may help the birds better cope with lead stress by actively maintaining protein structure, which may deform because of lead-related oxidative stress or lead replacing other metal ions in proteins.

As expected, the PP within-town comparison (PP High versus PP Low) showed substantially fewer significant proteins (Figure 7H). A total of 28 proteins were significantly differentially abundant, including BACH1 (upregulated in exposed birds) and members of the TRiC/CCT chaperonin complex (TCP1, CCT2, CCT3, CCT5, CCT6, CCT7, CCT8; all downregulated). Two proteins overlapped between the PP High versus PP Low comparison and the primary cohort: BACH1 and GPX1, with BACH1 increased, and GPX1 decreased in the PP exposure group (Figure 7I). Together, the recurrence of BACH1 differences between towns across independently prepared and analysed samples strengthens the evidence for this pathway’s role, though the unbalanced design and smaller sample sizes limit the strength of these comparisons.

### 3.10. Lead exposure induces hepatic metabolic reprogramming

A total of 3,358 protein groups were identified from the liver dataset, with a median coefficient of variation of 13.6% across pooled quality control samples. Because the hepatic proteome is significantly influenced by diet, dietary variation between populations could confound downstream proteome-based analyses of metabolic processes. The two towns compared in this study have similar climates and food availability, and dietary differences between bird populations are unlikely to account for observed proteomic changes. We first examined the within-town effect of lead toxicity by comparing birds with high blood lead concentrations to those with low concentrations within each site. Within Broken Hill, 86 proteins were differentially abundant between high-and low-lead groups (q-value < 0.05, log2FC > 0.5), of which 33 were increased in abundance in the exposure group (Figure 8a and Supplementary Table S12). The protein with the highest fold change in Broken Hill was metallothionein-2A (MT2A), a heavy metal detoxification and antioxidant defence protein, consistent with a direct molecular response to elevated lead burden [64, 65]. Conversely, the most significantly depleted protein was glutathione S-transferase alpha 1 (GSTA1), a key enzyme in glutathione-mediated detoxification, suggesting simultaneous induction of metal sequestration and erosion of antioxidant defence capacity under high lead exposure.

**Figure 8.**
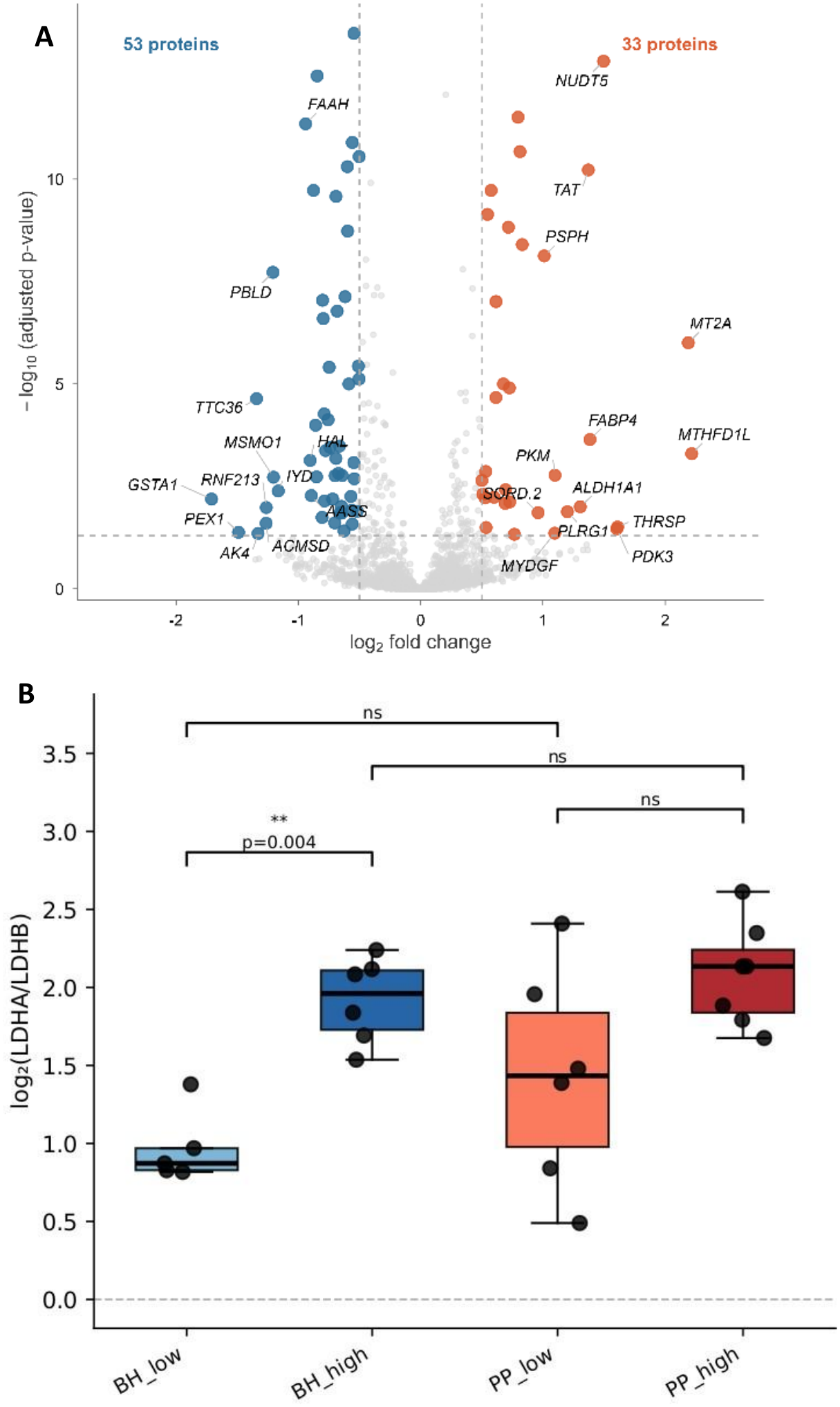
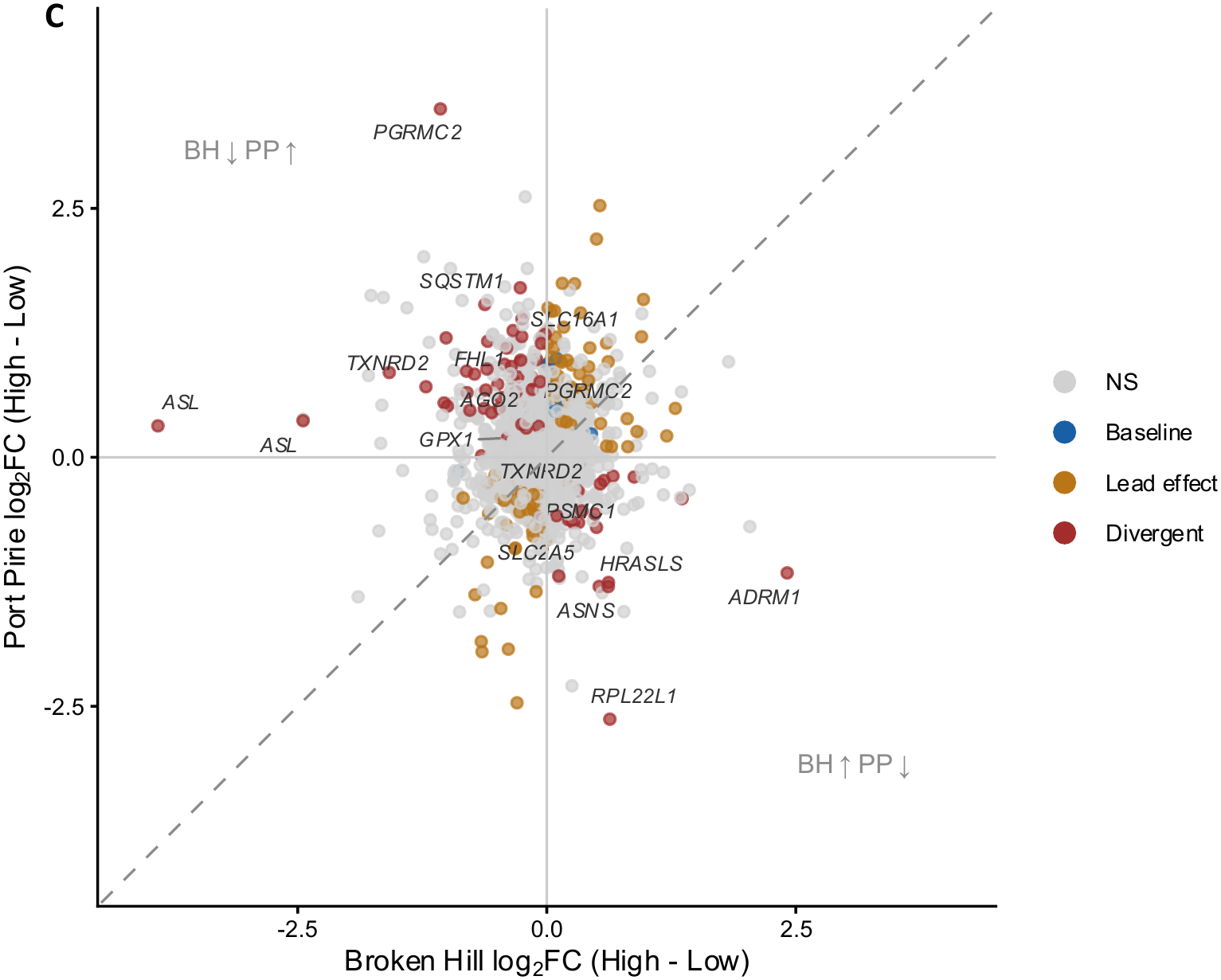
Liver proteomics reveals population-specific metabolic and proteostatic responses to lead. A) Volcano plot of the within-town lead effect in Broken Hill (BH High vs BH Low). B) Boxplots of lactate dehydrogenase A (LDHA) and lactate dehydrogenase B (LDHB) abundance in all four groups. C) Divergent response scatter plot showing per-protein lead response in Broken Hill (x-axis) versus Port Pirie (y-axis). Each point represents one protein; the dashed diagonal indicates identical response in both populations. Proteins in the off-diagonal quadrants respond to lead in opposite directions between the two populations.

Within Port Pirie sparrows, 125 proteins were significantly differentially abundant between the high- and low-lead groups, with 56 more abundant and 69 less abundant in the high-lead group (Supplementary Table S13). Gene Ontology (GO) enrichment analysis indicated that birds with high lead levels showed decreased amino acid catabolic processes across both sites, which is consistent with our cell line work (Figure. 7d). In Broken Hill, high lead birds also showed decreased abundance of enzymes involved in sterol biosynthesis, mitochondrial and peroxisomal beta oxidation (ACAD11, DBT, ABCD3, ACOX2, GNPAT, ACOX3), bile acid transport (ABCD3, ABCB11, ABCC2), and glutathione conjugation (GSTA1, MGST1, GSTK1), collectively indicating impaired amino acid and lipid metabolism and a reduced capacity for glutathione mediated detoxification of electrophilic toxins including lead. Compounding this deficit, the capacity to synthesise new glutathione may also be impaired, as glycine transporter 1 (SLC6A9), which imports glycine, the third substrate required for glutathione synthesis, was significantly decreased in both towns [66]. Multiple amino acid transporters were depleted in Port Pirie high lead birds, including SLC38A3, SLC7A2, SLC38A4, and SLC22A16, which suggests a broader suppression of amino acid import capacity at this site. KEGG pathway enrichment analysis of proteins decreased in abundance in high-lead birds identified valine, leucine, and isoleucine degradation and steroid biosynthesis as the most enriched pathways in Broken Hill, and glycine, serine, and threonine metabolism in Port Pirie (Supplementary Tables 14 and 15). For proteins more abundant in high-lead birds, the most enriched KEGG pathway in both towns was amino acid biosynthesis. This enrichment may reflect the transamination and redistribution of existing amino acid carbon skeletons toward metabolic priorities. This represents the single conserved hepatic stress response shared between the two sites.

### 3.11. Lead elicits a Warburg-like effect in the Broken Hill population

One of the key proteins differentially abundant in Broken Hill was lactate dehydrogenase A (LDHA), which catalyses the conversion of pyruvate to lactate in the final step of glycolysis, diverting pyruvate away from mitochondrial oxidation via the TCA cycle and toward anaerobic lactate production in the cytoplasm. Upregulation of LDHA has been associated with enhanced aerobic glycolysis in multiple cancer types [67, 68], and its increase in high lead Broken Hill birds suggests a shift toward glycolysis-dependent energy metabolism, consistent with the Warburg effect. This metabolic reprogramming is further supported by the LDHA-to-LDHB isoform ratio, in which LDHB catalyses the reverse reaction, converting lactate to pyruvate for entry into the TCA cycle. LDHB was significantly decreased in Broken Hill high lead birds, resulting in a significantly elevated LDHA to LDHB ratio in the high lead group relative to the low lead group (fold change 1.93, p < 0.01) (Figure. 8b). This isoform shift was absent in Port Pirie birds, where the ratio was unchanged between high- and low-lead groups and no changes were observed in HL-60 cell line treatment as well. The LDHA to LDHB isoform switch is further supported by the upregulation of pyruvate dehydrogenase kinase 3 (PDK3) in Broken Hill high-lead birds. PDK3 phosphorylates and inactivates the pyruvate dehydrogenase complex, thereby blocking the conversion of pyruvate to acetyl-CoA and preventing its entry into the mitochondrial TCA cycle [69]. Together, the LDHA-to-LDHB isoform shift and PDK3-mediated inhibition of pyruvate dehydrogenase constitute a coordinated mechanism that redirects pyruvate away from oxidative metabolism and toward cytoplasmic lactate production, collectively indicative of a Warburg-type metabolic phenotype in Broken Hill high-lead birds.

The sorbitol (polyol) pathway is a two-step metabolic pathway that converts glucose to fructose and has been directly implicated in the Warburg effect in cancer [70–72]. Fructose produced through this pathway bypasses the key regulatory enzymes of glycolysis, entering the glycolytic pathway as fructose-1-phosphate and generating unregulated flux that drives lactate production [70]. Fructose metabolism preferentially downregulates mitochondrial respiration while increasing aerobic glycolysis, and produces uric acid as a byproduct, which further inhibits the TCA cycle, generates mitochondrial ROS, and blocks fatty acid oxidation [70]. Both aldose reductase (AKR1B1) and sorbitol dehydrogenase (SORD) were significantly more abundant in Broken Hill high lead birds relative to Broken Hill low lead birds but maintained low levels in Port Pirie high lead birds. In the first step, aldose reductase reduces glucose to sorbitol, consuming NADPH as the hydride donor. In the second step, sorbitol dehydrogenase oxidises sorbitol to fructose using NAD+ as a cofactor and generating NADH as a byproduct [73]. The net biochemical consequence is a conversion of NADPH to NADH, consuming the primary cofactor required for antioxidant defence while generating a reducing equivalent that accumulates under conditions of mitochondrial dysfunction. Glutathione reductase requires NADPH to recycle oxidised glutathione back to its active reduced form, and polyol pathway activation has been shown to decrease both glutathione and NADPH levels by reducing NADPH availability for glutathione reductase [74]. The NADH generated by sorbitol dehydrogenase further inhibits the TCA cycle and pyruvate dehydrogenase complex, driving additional glucose flux into the polyol pathway.

This NADH accumulation constitutes a state of reductive stress, which is further exacerbated by the concurrent decreased abundance of NDUFC2, a structural subunit of Complex I, thereby impairing the primary mechanism for mitochondrial NADH oxidation and NAD+ regeneration. With Complex I unable to efficiently process NADH, and the polyol pathway simultaneously generating additional NADH while consuming NADPH, Broken Hill high-lead birds face a compounding redox crisis in which insufficient NADPH for antioxidant defence and accumulating NADH drive progressive ROS generation. The upregulation of LDHA under these conditions therefore serves not only as a marker of metabolic reprogramming but also as an active mechanism to relieve reductive stress and regenerate cytoplasmic NAD+ from accumulating NADH, thereby partially offsetting the mitochondrial bottleneck imposed by Complex I impairment.

The Warburg effect is canonically driven by hypoxia-inducible factor 1 (HIF-1), which mediates a transcriptional programme that suppresses pyruvate entry into the TCA cycle and inhibits fatty acid beta oxidation, another major source of acetyl-CoA [75]. The suppression of both mitochondrial and peroxisomal beta-oxidation enzymes observed in Broken Hill high-lead birds is consistent with this HIF-1-mediated metabolic programme [76]. Although HIF-1 itself was not detected in the liver or HL-60 cell line datasets, the coordinated proteomic changes observed in Broken Hill high-lead birds, encompassing glycolytic commitment, blockade of pyruvate oxidation, and suppression of beta-oxidation, are consistent with activation of a HIF-1-like transcriptional response under conditions of lead-induced mitochondrial dysfunction.

### 3.12. In exposed birds, major differences lie in the mitochondrial proteome

The high lead level groups from each town were compared to identify which proteins were differentially abundant between them. We identified 98 differentially abundant proteins, with 33 more abundant in Broken Hill birds and 65 in Port Pirie birds (Supplementary Table 16). More than 50% of the differentially abundant proteins are mitochondrial proteins. GO enrichment analysis indicated that Port Pirie is enriched in oxidative phosphorylation and glutathione metabolism. In Broken Hill high lead birds, the top enriched KEGG pathway was glycine, serine, and threonine metabolism. Importantly, when low lead birds from both towns were compared, Broken Hill low lead birds were also enriched in the glycine, serine and threonine metabolism pathway relative to Port Pirie, but in this case, both catabolic and anabolic enzymes of the pathway were elevated simultaneously.

An interaction model was fitted to the liver proteomics data and identified 110 proteins whose lead response differed significantly between populations (p < 0.05, |interaction log2FC| > 0.25; Figure 8C). On the Port Pirie side, PGRMC2 (progesterone receptor membrane component 2) and SQSTM1 (sequestosome 1/p62) were the most prominent divergent proteins. PGRMC2 showed one of the largest interaction effects in the dataset, with PP birds strongly upregulated under lead (+3.50 log2FC), while BH birds showed a slight decrease (−1.07 log2FC; interaction p = 0.020). PGRMC2 functions as an intracellular haem chaperone that transports signalling haem from the mitochondria to the nucleus, where haem serves as a regulatory ligand for the transcription factor BACH1. Under normal conditions, haem binding promotes BACH1 nuclear export and proteasomal degradation, thereby relieving repression of haem-responsive and antioxidant genes. Although BACH1 was not reliably quantified in the liver data, the blood proteomics demonstrated that BACH1 accumulates in PP lead-exposed birds because of ALAD inhibition and haem depletion. The increased abundance of PGRMC2 in PP livers may therefore represent a compensatory response to increase nuclear haem delivery and promote BACH1 degradation, effectively attempting to restore antioxidant gene expression through enhanced haem trafficking rather than by maintaining haem synthesis at the ALAD level. ALAD itself was not differentially abundant in the liver (interaction p = 0.499). SQSTM1/p62, the canonical selective autophagy receptor, showed a striking liver-specific divergence: PP birds strongly increased it under lead (+1.54 log2FC) while BH birds suppressed it (−0.62 log2FC, interaction p = 0.023). This contrasts with the blood, where SQSTM1 was increased in abundance in both populations to a similar extent (BH +0.90, PP +0.76, interaction p = 0.83). The liver-specific divergence suggests that PP birds deploy selective autophagy as a primary proteostasis mechanism in the liver, consistent with the broader pattern of PP favouring autophagic clearance over proteasomal degradation.

In the Broken Hill population, ADRM1 (adhesion regulating molecule 1/proteasomal ubiquitin receptor) showed the most extreme divergence, increasing dramatically in BH (+2.41 log2FC) while decreasing in PP (−1.16 log2FC, interaction p = 0.012). ADRM1 sits on the 19S regulatory particle of the 26S proteasome, and recruits ubiquitinated substrates for degradation. Its increased abundance in BH livers, alongside a significant interaction with PSMC1 (an AAA-ATPase subunit of the 26S proteasome; p = 0.006), indicates that BH birds actively enhance proteasomal substrate delivery and processing capacity under lead stress. This mirrors the proteasome upregulation observed in BH blood and reinforces the cross-tissue pattern that BH populations favour the ubiquitin-proteasome pathway.

## 4. Conclusions and Limitations

In this study, we performed proteomic analysis of blood and liver tissue from house sparrows (*Passer domesticus*) inhabiting two lead-contaminated mining towns in Australia. Broken Hill represents a population that has been geographically isolated for over a century, while Port Pirie maintains substantial gene flow with surrounding non-contaminated populations. Our results reveal proteomic divergence between these two populations, both at baseline and in their molecular responses to chronic lead exposure. Despite the limited sample sizes inherent to field-based wildlife proteomics, we demonstrate that Broken Hill birds exhibit a distinct proteomic profile that diverges considerably from Port Pirie. However, this divergence does not necessarily reflect positive adaptation. ALAD provides a clear example: while the protein itself showed significant population-level divergence, elevated ALA concentrations in Broken Hill birds indicate that the haem biosynthesis pathway remains functionally disrupted, suggesting that the haemoglobin-related phenotypic differences may have arisen through compensatory mechanisms in alternative pathways rather than through restored ALAD function. Additionally, the upregulation of proteasomal and ubiquitin-dependent degradation pathways in Broken Hill birds may reflect an increased reliance on protein quality control systems to maintain structural integrity under chronic lead exposure. To determine whether lead-induced protein misfolding is the underlying driver of this response, we propose applying limited proteolysis-coupled mass spectrometry (LiP-MS) to quantify proteome-wide structural alterations across populations. Critically, the present dataset cannot resolve whether the observed proteomic differences arise from phenotypic plasticity in response to local lead exposure or from heritable genomic variation fixed through generations of selection. Distinguishing between these mechanisms requires multi-omics integration across genomic, transcriptomic, and proteomic layers. Whole-genome sequencing data from both populations is available and will be incorporated in subsequent analyses alongside planned transcriptomic profiling, to determine the extent to which the molecular divergence observed here is encoded in the genome and how these changes collectively shape organismal responses to chronic heavy metal stress.

## 5. Acknowledgments

Some aspects of the research described herein were facilitated by access to the Australian Proteome Analysis Facility (APAF) funded under the Australian Government National Collaborative Research Infrastructure Strategy (NCRIS)/Education Investment Fund.

## Notes

### Competing Interest Statement

The authors have declared no competing interest.

